# Gut Microbiota-Targeted Diets Modulate Human Immune Status

**DOI:** 10.1101/2020.09.30.321448

**Authors:** Hannah C. Wastyk, Gabriela K Fragiadakis, Dalia Perelman, Dylan Dahan, Bryan D Merrill, Feiqiao B. Yu, Madeline Topf, Carlos G. Gonzalez, Jennifer L. Robinson, Joshua E. Elias, Erica D. Sonnenburg, Christopher D. Gardner, Justin L. Sonnenburg

## Abstract

Diet modulates the gut microbiome, and gut microbes, in turn, can impact the immune system. Here, we used two gut microbiota-targeted dietary interventions, plant-based fiber or fermented foods, to determine how each influences the human microbiome and immune system in healthy adults. Using a 17-week randomized, prospective study design combined with -omics measurements of microbiome and host, including extensive immune profiling, we found distinct effects of each diet. High-fiber consumers showed increased gut microbiome-encoded glycan-degrading CAZymes despite stable community diversity. Three distinct immunological trajectories in high fiber-consumers corresponded to baseline microbiota diversity. Alternatively, the high-fermented food diet steadily increased microbiota diversity and decreased inflammatory markers. The data highlight how coupling dietary interventions to deep and longitudinal immune and microbiome profiling can provide individualized and population-wide insight. Our results indicate that fermented foods may be valuable in countering the decreased microbiome diversity and increased inflammation pervasive in the industrialized society.

## Introduction

The importance of the gut microbiota, or microbiome, in human health (Lynch and Pedersen, 2016) necessitates enhanced understanding of the factors that influence the composition and function of this microbial community. Diet has emerged as a driving factor in microbiota composition and function (Flint et al., 2017; Muegge et al., 2011; Rothschild et al., 2018; Zhernakova et al., 2016). The profound link between diet and the microbiota in humans has been demonstrated in numerous ways, for example, by coupling long-term dietary patterns and microbiota diversity, taxonomic composition, and microbiome gene content measurements (Smits et al., 2017; Jha et al., 2018; Yatsunenko et al., 2012; Arumugam et al., 2011). Short-term changes in diet during prospective dietary intervention studies have also been shown to rapidly change the human gut microbiota (David et al., 2014). Although, we and others have reported a general resilience of the human microbiota over short time periods (days to months) coupled with retention of highly individualized microbiome identities (Wu et al., 2011; Fragiadakis et al., 2020).

The integration of the gut microbiota into human biology suggests that manipulation of gut microbes may be a powerful means to alter diverse aspects of human health. Diets targeting the gut microbiome to enhance, introduce, or eliminate specific functionalities or taxa, could prove a powerful avenue for realizing the promises of precision medicine. Even in the absence of manipulation, the gut microbiome contains features that are informative when predicting individual-specific postprandial responses to specific foods (Zeevi et al., 2015; Brand-Miller and Buyken, 2020). One key question is whether there are broad, non-personalized dietary recommendations that can leverage extant microbiota-host interactions for improved health across populations.

Non-communicable chronic diseases (NCCDs) are largely driven by chronic inflammation and rates are increasing rapidly with industrialization. Coincidentally, gut microbiota changes with industrialization are also well-documented. Rapid “westernization” of the microbiota has been observed in U.S. immigrants, with loss of microbial functions and taxa accompanied by deteriorating markers of host health, increased BMI, and rising inflammatory markers typical in industrialized populations (Vangay et al., 2018; Sonnenburg and Sonnenburg, 2019). A 2-week food exchange study in which African Americans consumed a rural African diet and rural Africans ate a typical African American diet revealed measurable changes to the microbiota and markers of cancer risk despite the brevity of the dietary intervention (O’Keefe et al., 2015). Given that the human microbiome is known to influence inflammatory status, a key question is whether diets that target the gut microbiome can attenuate systemic inflammation in healthy individuals.

A diverse body of literature supports the role of fiber in health including a dose-response relationship of higher fiber consumption and lower rates of mortality (Liu et al., 2015). Mechanistic studies in animal models reveal the role of microbiota accessible carbohydrates (MACs) present in dietary fiber in supporting gut microbiota diversity and metabolism, and the positive role of short-chain fatty acids, a product of fiber fermentation by the gut microbiota, in maintaining gut barrier health and attenuating inflammation (reviewed in Makki et al., 2018; Sonnenburg and Sonnenburg, 2014). Dietary interventions that specifically alter dietary fiber, such as increases in total carbohydrates, whole grains, and resistant starch versus wheat bran consumption, have shown impacts on the microbiota along with improvements in health markers of the study participants (Duncan et al., 2007; Martínez et al., 2013; Walker et al., 2011). These findings and the shortfall between fiber consumption in the average American diet versus recommended levels suggest that boosting fiber intake could be a powerful way to modulate the human immune system via the microbiota (Deehan and Walter, 2016).

Fermented foods, such as kombucha, yogurt, and kimchi, have gained popularity as reports of potential health benefits in animal models and humans have emerged (Dimidi et al., 2019). Large cohort studies as well as limited interventional studies have linked the consumption of fermented foods with weight maintenance and decreased diabetes, cancer, and cardiovascular disease risks (Mozaffarian et al., 2011; Díaz-López et al., 2016; Gille et al., 2018). A recent longitudinal study of a subset of American Gut Project participants found differences in microbiota composition and fecal metabolome among fermented food consumers vs. non-consumers (Taylor et al., 2020). Given that fermented foods have historically been part of many diets around the world, consuming fermented foods may offer an effective way to reintroduce evolutionarily important interactions. They may also provide compensatory exposure to safe environmental and foodborne microbes that have been lost over the course of sanitizing the industrialized environment.

To address whether microbiota-targeted diets can positively impact human biology, we have performed a dietary intervention while longitudinally monitoring the microbiome and immune status in healthy adults. Here we study the effect of two diets, high fermented or high fiber food (Fermented and Fiber-rich Food (FeFiFo) Study; ClinicalTrials.gov Identifier: NCT03275662), on the human immune system using -omics profiling including state-of-the-art immune profiling, in a randomized, prospective study design. We observed that each intervention produced a distinct response and that some responses were general, i.e., cohort-wide, while others were individualized. Remarkably, over the course of the 10-week intervention, we observed a cohort-wide decrease in many inflammatory markers in individuals consuming fermented foods, coincident with an increase in microbiota diversity. These results suggest that fermented foods may be powerful modulators of the human microbiome-immune system axis and may provide an avenue to combat NCCDs.

## Results

### Participants successfully increased their assigned dietary fiber or fermented food consumption over the course of the study

In order to examine the effect of diet on the microbiome and the immune system, generally healthy adults were recruited to participate in a 10-week dietary intervention (17-week protocol including pre- and post-intervention) in which participants were randomized to one of two diet arms: a high-fiber diet or a high-fermented foods diet (Figure 1A, Table S1). Of 381 individuals assessed for eligibility, 39 participants were assigned (37 randomized (R), 2 non-randomized (NR)) to one of the two interventions: a high-fiber diet (n=21, 19 randomized (R), 2 non-randomized (NR)) or a high-fermented foods diet (n=18). One participant dropped out of the study due to personal reasons and two participants were prescribed antibiotics during the course of the study and were excluded from analysis. The final count for participants was identical in each arm, n=18. Participants were adults (age 51 ± 12 y [mean ± SD]), with a mean BMI of 25 ± 4 kg/m2, predominantly women (73%) and Caucasian (81%), and with a high education level (89% with a college degree or higher) (Table S1). The study was approved annually by the Stanford University Human Subjects Committee. Blood and stool samples were collected longitudinally along a 3-week pre-intervention time period (“baseline”), followed by a 4-week ramp phase where participants gradually increased intake of their respective diets (“ramp”), then a 6-week maintenance phase where participants maintained a high-level of consumption either fiber or fermented foods (“maintenance”), and finally a 4-week choice period where participants could maintain their diet to their desired extent (“choice”, Figure 1B). Stool samples were assessed for microbiota composition, function, and metabolic output. Blood samples were used to generate a systems-level view of the immune system including measurements of circulating cytokine levels, cell-specific cytokine response signaling, and cell frequency and immune cell signaling at steady-state (Figure 1B, STAR methods). The number of participant samples analyzed for each experimental platform and time point varied slightly depending on sample availability (Table S2). Importantly, participants’ gut microbiota at baseline did not differ between the two arms, as determined by alpha and beta diversity measurements (data not shown).

**Figure 1.**
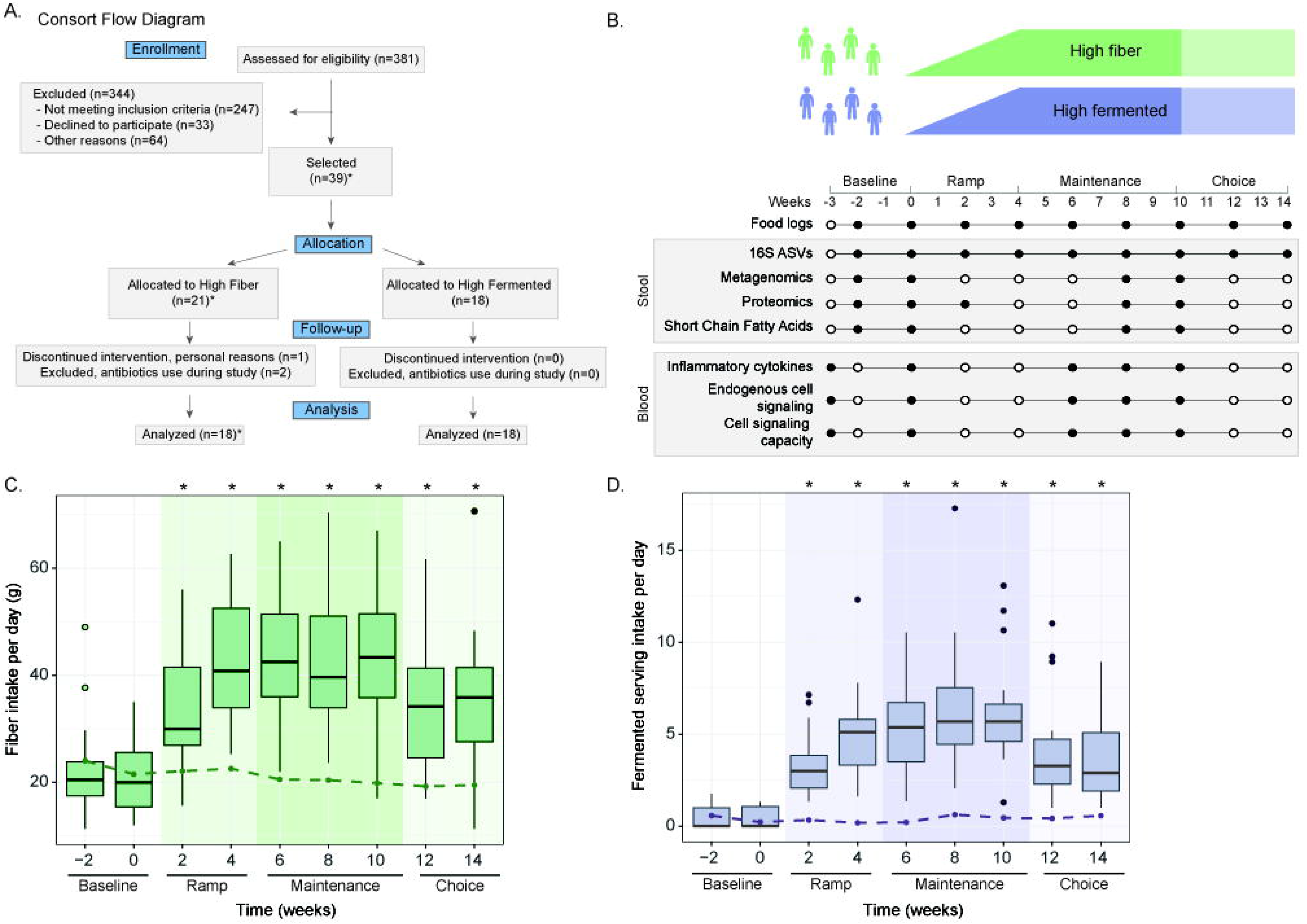
Overview of Fiber Fermented Food Study. **(A)** Consort flow diagram for participant enrollment, allocation, follow-up, and analysis. Side chart shows the number of participants in high-fiber (Fi) and high-fermented (Fe) diet arm collected for each platform. *2 participants were assigned to high fiber, not randomized, by special request. **(B)** The 14-week study overview timeline, sample types collection, and corresponding experimental platforms. **(C)** Fiber intake in the high-fiber diet arm shown in boxplots, fiber intake in the high-fermented food diet arm shown as dotted line. **(D)** Fermented food intake in the high-fermented food diet arm shown in boxplots, fermented food intake in high-fiber diet arm shown as dotted line. P-values <0.05 via t-test denoted by asterisks and calculated for each time point relative to baseline -2 week value.

Participants successfully increased their consumption of fiber or fermented foods as determined by macronutrient and micronutrient data extracted from food logs generated by participants every two weeks. Those in the high-fiber diet arm increased their fiber consumption from an average of 21.5±8.0 g per day at baseline to 45.1±10.7 g per day at the end of the maintenance phase (Figure 1C). Participants in the high-fermented food diet arm consumed an average of 0.4±0.6 servings per day of fermented food at baseline, which increased to an average of 6.3±2.9 servings per day at the end of the maintenance phase (Figure 1D). Importantly, participants in the high-fiber diet arm did not increase their consumption of fermented foods (Figure 1C, dashed line), nor did participants consuming the high-fermented food diet increase their fiber intake (Figure 1D, dashed line) during the course of the study.

An analysis of selected macro- and micronutrients revealed differences from baseline to the end of the maintenance phase in the consumption of several nutrients in the high-fiber arm. High-fiber diet participants increased their intake of soluble and insoluble fiber, carbohydrates, vegetable protein, and had a modest increase in calories, along with increases in iron, magnesium, potassium, vitamin C, and calcium. These participants also decreased their consumption of animal protein and sodium (Table S3). Conversely, the high-fermented food diet participants increased their intake in animal protein due to the increased consumption of fermented dairy products. Notably, despite higher consumption of fermented vegetables and vegetable brine drinks, total sodium intake did not change in the fermented food arm compared to their baseline diet (Table S3). In order to gain a more detailed understanding of how participants increased their fiber or fermented foods intake, fiber-rich and fermented foods were grouped into subcategories (see STAR methods). Fiber-rich foods were categorized into fruits, vegetables, legumes, grains, nuts and seeds, and other. Fermented foods were grouped into yogurt, kefir, fermented cottage cheese, fermented vegetables, vegetable brine drinks, kombucha, other fermented non-alcoholic drinks, and other foods. While all participants followed the requirements for the dietary intervention, each implemented the intervention differently in terms of specific subcategories of fiber-rich or fermented foods they consumed (Figure S1A, S1B).

The primary outcome of cytokine response score difference from baseline to end of intervention was not significant for either arm of the study. However, several changes in secondary and exploratory outcomes were observed. Decreases in inflammatory markers and increases in microbiota diversity from baseline to end of intervention were significant in the fermented food arm, a specific subset of short chain fatty acids (SCFAs) were significantly decreased in the high-fiber arm from baseline to end of intervention. These results are discussed in greater detail later in the manuscript. For a full report of primary and secondary outcome results please see Table S4. To assess participants overall health throughout the study, blood glucose, insulin, triglycerides, LDL-C, HDL-C, blood pressure, and waist circumference were measured; however, no differences were observed in this generally healthy cohort either between the two arms or longitudinally (within) (data not shown). Based on assessment from the Gastrointestinal Symptoms Rating Scale (Svedlund et al., 1988) participants on the high-fiber diet reported an increase in stool softness from baseline (average Bristol stool type = 3.3 +/− 0.3) to end of ramp phase (stool type=4.1 +/− 0.3; p=0.04; paired t-test) and the end of maintenance phase (stool type=4.1 +/− 0.3, p=0.004; paired t-test). Whereas, the fermented food diet arm reported an increase in bloating from baseline (abdominal distention score=0.06 +/− 0.06) to the end of the ramp phase (score=0.4 +/− 0.1; p=0.03; paired t-test), which was no longer significant by the end of the maintenance phase. Additional validated surveys were given to participants to assess perceived stress, well-being, fatigue, physical activity, and cognition; however, no significant changes were observed between the two groups or longitudinally within the groups (data not shown).

### A high-fiber diet and a high fermented foods diet result in distinct effects on the gut microbiota and host immune system

Given the success that participants achieved in adhering to their assigned dietary intervention, we wondered whether each diet intervention produced a characteristic change in participants’ microbiota or aspects if their biology. We generated random forest models in which a separate model was made for each assay (listed in Figure 1B), using features quantified as the difference between baseline (week -2 for stool samples, week -3 for blood samples) and the end of the maintenance phase (Week 10) (Figure 2A). Each model used recursive feature elimination to select models with the least number of features while maintaining the highest accuracy. (Features identified for each model are listed in Table S5.) As a positive control, nutritional intake was used in its own model, which classified participants by diet arm with 91% accuracy (leave one out cross-validation, LOOCV). This model relied upon intake of animal protein, total dietary fiber, and insoluble dietary fiber for classification. As a negative control, the two baseline time points were used as parameters for the model and had a prediction accuracy equivalent to chance (48%).

**Figure 2.**
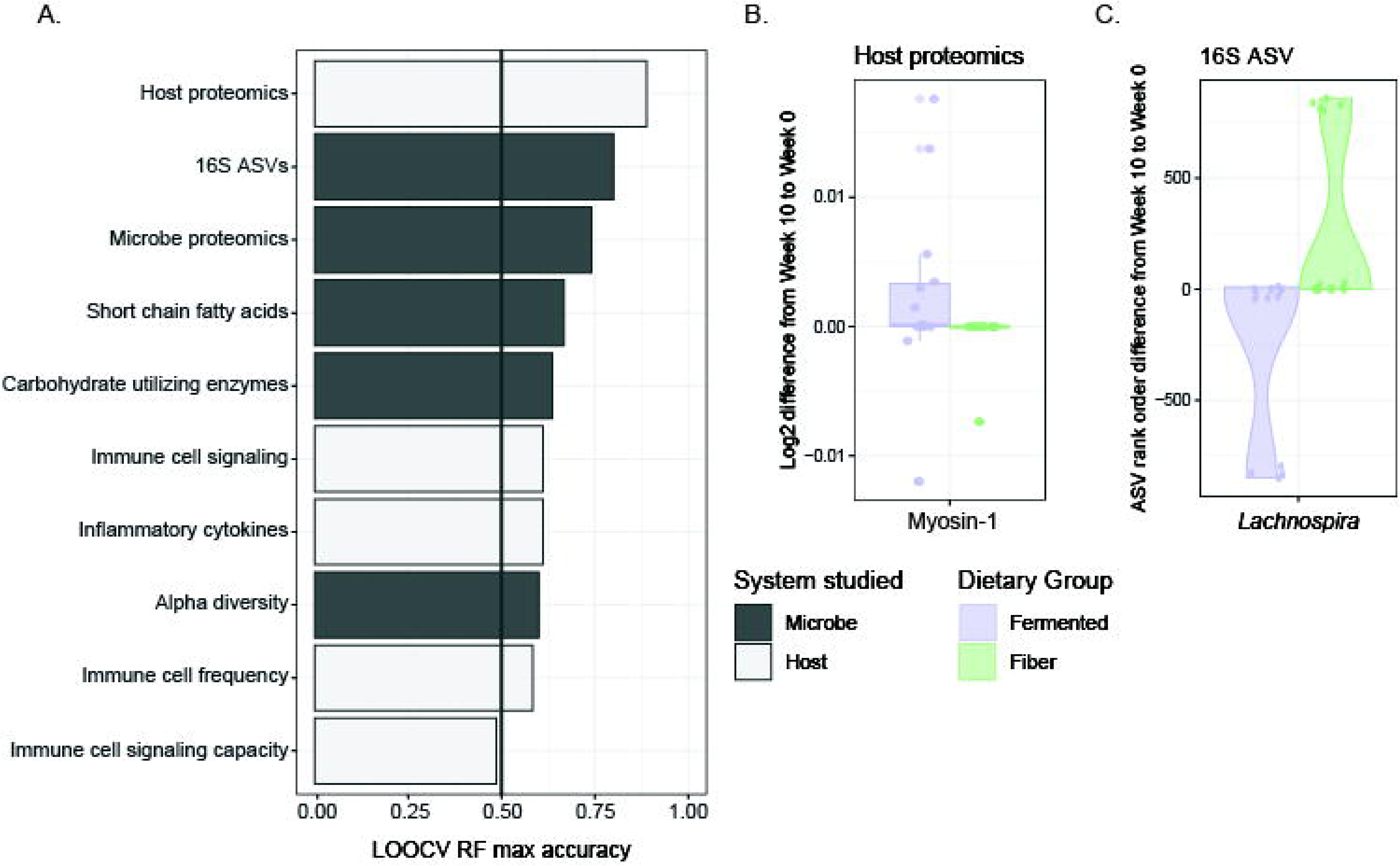
Diet-specific effects of a fiber vs. fermented food intervention on the host and microbiome. **(A)** Accuracy of leave-one-out cross-validation (LOOCV) of random forest models predicting diet group; separate models using host-derived data (white bars) or microbe-derived data (black bars), using parameter changes from baseline to end of maintenance as model features. Recursive feature elimination chose the minimum number of parameters needed for maximum accuracy. **(B)** Differences in myosin-1, model feature selected for host proteomics model. **(C)** Differences in rank-order change of Lachnospira, model feature selected for 16S amplicon sequence variants (ASVs) model. Purple, fermented group; green, fiber group.

Aside from nutrition intake, the highest accuracy in predicting diet was achieved with the model that used human stool proteomic changes (89% accuracy). This model selected a single parameter from the 230 input host proteins, myosin-1, which increased in the high-fermented food diet arm (Figure 2B). Myosin-1 is highly expressed in microvilli within the brush border of the small intestine. Its increase could indicate increased epithelial cell turnover within the small intestine in the high-fermented food diet cohort (Benesh et al., 2010). A more detailed analysis on the proteomic differences between the high-fiber and high-fermented food diet arms has been reported previously (Gonzalez et al., 2020).

The four next highest-performing models were all generated using measurements of the microbiota including overall 16S rRNA-based composition (80% accuracy), stool microbe proteomics (74%), stool short-chained fatty acids (67%), and metagenomic measurement of carbohydrate active enzymes (64%). Microbiota diversity was a less effective model (60%) in discerning which diet a participant followed. The overall microbiota composition model was characterized by an increase in the genus *Lachnospira* in the high-fiber diet arm and a decrease in the high-fermented food arm relative to baseline (Figure 2C). *Lachnospira* has been positively associated with high dietary fiber consumption in a human prospective study (Lin et al., 2018). We found that in both diet arms participants’ gut microbiota composition stayed highly individualized during the intervention rather than clustering by diet at the end of the intervention, findings similar to those reported in previous human microbiota studies (Johnson et al., 2019; Wu et al., 2011). However, using Bray-Curtis beta diversity, the linear regression of the distance from centroid versus time using a linear mixed effects model had a negative coefficient for the both the high-fiber and high-fermented food diet arms (slope = −4.2e-3, p-value = 1.6e-3; and slope = −5.3e-3, p-value = 1.4e-4, respectively). In other words, an individual’s microbiota composition became more similar to that of other participants within the same arm over the intervention, despite retaining the strong signal of individuality.

Models generated using measurements of host immune parameters including endogenous immune cell signaling (61%), inflammatory cytokines (61%), and immune cell frequency (58%) were predictive of diet arm, but less so than many of the observed changes to the microbiota listed above. Models generated using immune cell signaling capacity (49%) were no better than chance at predicting diet group. These models demonstrate that the two dietary interventions produced characteristic responses in participants’ human and microbial biology. Diet-induced changes to the microbiota may be more consistent across individuals than immune system responses, in accordance with the interventions targeting the gut microbiota.

### Fiber intake shifts carbohydrate processing capacity and metabolic output of the microbiota

Since predictive models revealed diet-specific responses in participants, we investigated more thoroughly how each diet impacted the host microbiota. The high-fiber diet arm was assessed for changes in microbiota composition, diversity, function, and microbially-derived products of fermentation. Based on interventional studies in mice and humans and long-term association studies between high-fiber diets and increased microbiota diversity, we hypothesized that increasing dietary fiber consumption would lead to an increase in microbiota diversity (Sonnenburg et al., 2016; De Filippo et al., 2010; Cotillard et al., 2013; Le Chatelier et al., 2013). However, alpha diversity, as determined by the number of observed ASVs, Shannon diversity, or phylogenetic diversity, did not change cohort-wide over the course of the intervention (Figure 3A, Figure S2). Nor were there changes in alpha diversity when correlated with the overall quantity of dietary fiber consumed per participant as determined using a linear mixed effects model varying fiber intake versus alpha diversity across study and correcting for participant (data not shown). However, we did observe an increase in the relative abundance of microbial proteins per gram stool of the high-fiber diet arm from baseline (Week -2 and Week 0) to the end of the maintenance phase (Week 10), suggesting that the density of microbes within the microbiota may have increased with higher fiber consumption (Figure 3B). No specific taxon (ASV) exhibited altered relative abundance over time across the entire high-fiber diet arm. While increased *Lachnospira* relative abundance was observed in the high-fiber diet arm when compared to the high-fermented food diet arm (Figure 2C), this genus was not significantly increased within the high-fiber diet arm from baseline to the end of the maintenance phase. This discrepancy could be due to the lower sample size when comparing within one diet arm alone or due to decreasing relative abundance of *Lachnospira* in the high-fermented food diet arm over the course of the intervention.

**Figure 3.**
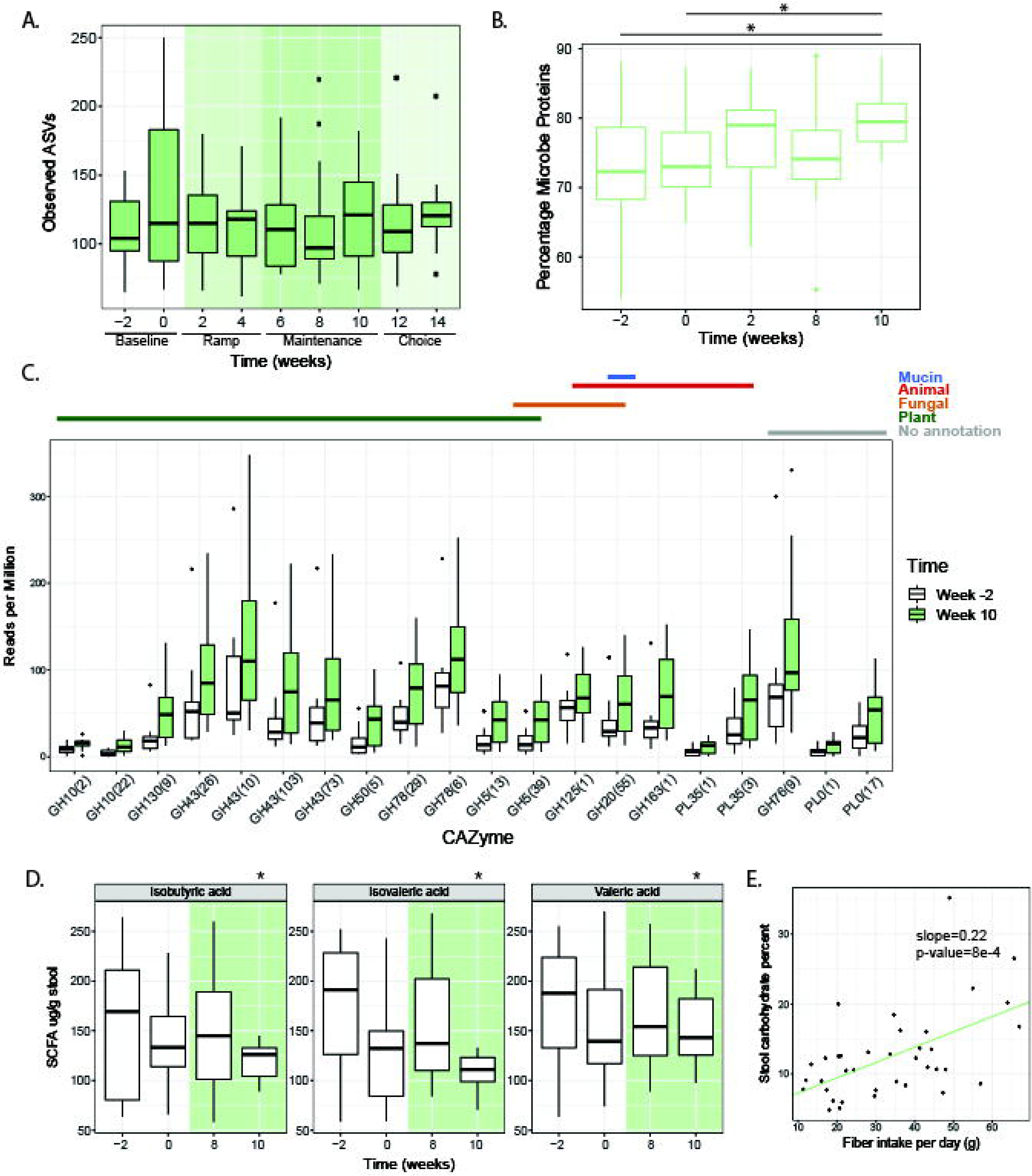
Participants consuming fiber exhibit shifts in the functional profile of the microbiome. **(A)** Observed number of amplicon sequence variants (ASVs) from 16S rRNA amplicon sequencing; no significant changes during any intervention time point compared to baseline (week -2 or Week 0) (paired t-test). **(B)** Proteins measured using LC-MS (Gonzalez et al., 2020) were categorized as human or microbe derived using the HMP1 database (The Human Microbiome Project Consortium, 2012). Microbe proteins as a percent of total stool proteins increase from baseline to end of maintenance phase (Week 10, p-value=0.003 from week -2, p-value=0.01 from Week 0, paired t-test). **(C)** CAZymes identified from metagenomic sequencing as significantly changing in relative abundance from baseline to end of maintenance phase (FDR ≤ 0.05, q-value ≤ 0.1, SAM two-class paired). CAZymes were annotated using dbCan and assigned to functional categories (Yin et al., 2012; Cantarel et al., 2012). **(D)** Significant decreases in two branched chain fatty acids and valeric acid in stool (p-value=0.044, 0.033, 0.033, paired t-test). **(E)** Total fiber intake (grams) correlated with percentage of carbohydrates in stool using linear mixed effects (LME) model (p-value=8e-4).

Despite the lack of generalized changes to microbiota diversity or composition in the high-fiber diet arm, the increased percentage of microbial proteins in the stool suggests that fiber may fuel the growth of bacteria adept at fiber-degradation. Metagenomic sequencing revealed an increase in the relative abundance of 21 different carbohydrate active enzymes (CAZymes); of 15 putative non-ambiguous substrate predictions, ten are predicted to degrade plant cell wall carbohydrates (Figure 3C). There were no CAZymes that decreased in relative abundance, indicating that the high-fiber diet led to an overall increase in complex carbohydrate processing capacity cohort-wide and not just a reconfiguration of carbohydrate utilization functionality. Therefore, while fiber consumption appears to consistently increase CAZyme abundance, the taxonomic changes that result in these increases may differ across participants. The individualized taxonomic solutions to increasing CAZymes may reflect the individualized collection of microbes in each person’s gut but may also be a result of the different types of complex carbohydrates consumed by the participants who were eating non-identical high-fiber diets.

To assess the metabolic output of the microbiota in the high-fiber diet arm, levels of fecal short-chain fatty acids (SCFAs) were measured. We did not observe an increase in butyrate as has been previously reported in some studies of dietary fiber consumption (So et al., 2018). However, significant heterogeneity exists in butyrate results between dietary fiber studies, possibly due to incomplete fermentation and/or colonic absorption by the host (So et al., 2018). We did observe a decrease in the branched-chain fatty acids (BCFAs) isobutyric and isovaleric acid, as well as valeric acid from baseline to the end of the maintenance phase (Figure 3D). Elevated isobutyric and isovaleric acid have been associated with hypercholesterolemia (Granado-Serrano et al., 2019) and elevated valeric acid in autism spectrum disorder (Liu et al., 2019). It is not clear whether these changes in BCFAs are a result of decreased production by the microbiota or decreased consumption of dairy and beef, which contain high levels of BCFAs (Ran-Ressler et al., 2014).

Despite observed changes in CAZyme profiles and SCFA levels, we were surprised that given the substantial increase in fiber consumption in this arm, there was not a larger microbiota response. We wondered whether the intervention was too brief for the microbiota to adequately adapt to the increase in fiber consumption. Specifically, we wondered whether increased consumption of fiber was overwhelming the fermentative capacity of the participants’ microbiota. Carbohydrates were extracted from stool samples, acid-hydrolyzed to release monosaccharides, and then measured using HPLC. A significant correlation was observed between increased fiber consumption among participants and an increase in total stool carbohydrates (p-value=8e-4; LME correcting for participants across time, Figure 3E). These data suggest the carbohydrate degradation by participants’ microbiota was insufficient to process the increased fiber consumption, consistent with analyses of industrialized microbiome (Smits et al., 2017; Vangay et al., 2018). It is possible that a longer intervention would have allowed for adequate microbiota remodeling and recruitment from external sources. Alternatively, the deliberate introduction of fiber-consuming microbes may be required to increase the microbiota’s fermentative capacity.

### High-dimensional immune system profiling reveals sub-types of host responses to fiber intake

Changes to the microbiota in the high-fiber diet arm led us to wonder whether participants’ immune system was coincidentally impacted. We tested whether participants’ immune status, as measured by assays selected to capture complementary aspects of immune cell signaling activity both intracellularly and through cytokine mediators, was altered (Figure 4A). A multiplex proteomic platform assessed circulating cytokines and additional immune modulators in serum with a panel specific for inflammation (Olink technology, Table S6). Whole blood was subjected to single-cell mass cytometry (CyTOF, Table S6, Figure S3) with a 50-parameter panel of antibodies to delineate major immune cell types (cell frequencies) and activation of canonical immune cell signaling pathways (endogenous signaling, Table S6). Finally, immune signaling capacity was measured by stimulating peripheral blood *ex vivo* with lipopolysaccharide (LPS) or one of five cytokines involved in inflammatory signaling (IL-6, IL-2, IL-10, IFNa, IFNg); cell type-specific responses were measured in the JAK/STAT and MAP kinase pathways by flow cytometry (Table S6). From each of these assays, a set of immune features was derived as a descriptor of immune activity (see STAR Methods).

**Figure 4.**
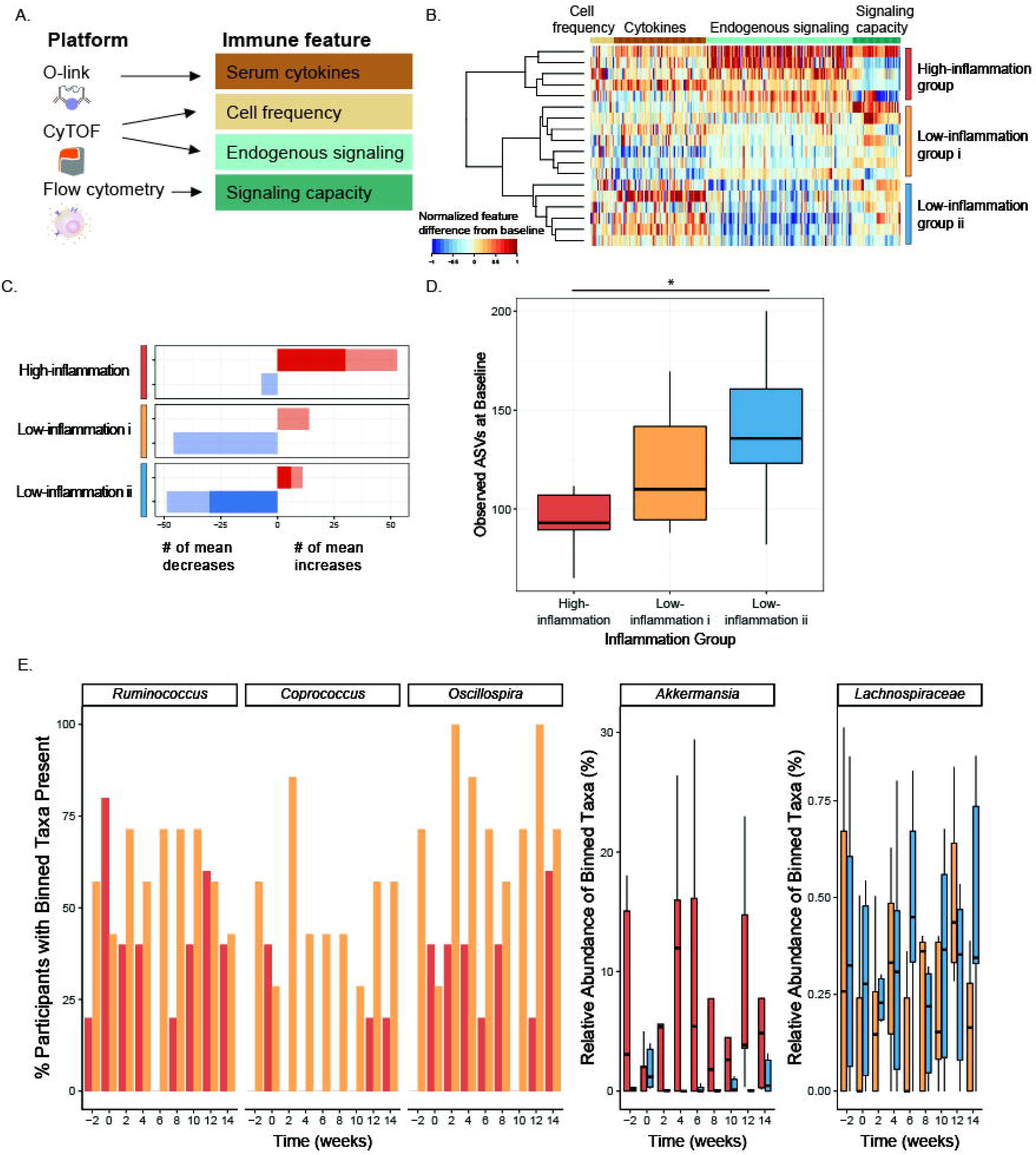
Fiber-consuming participants exhibit varied immune responses that track with differences in microbiome composition and diversity. **(A)** Immune features derived from immunophenotyping assays. **(B)** Heatmap depicting differences in immune features (grouped by feature type) from baseline (week -3) to end of intervention (Week 10), rescaled from minimum change > −1 to maximum change < 1, no change=0. Each row is a participant, rows are clustered using hierarchical clustering by feature values in the fiber arm. **(C)** Counts of the mean positive (red) or mean negative (blue) changes in endogenous immune cell signaling from baseline (week -3) to end of maintenance (Week 10) for the three clusters. Non-significant changes shown in light color, significant changes shown in dark color (SAM, two-class paired, q-value<0.1). **(D)** Average number of observed ASVs at baseline (week -2 and Week 0) high-inflammation cluster (red), low-inflammation i cluster (gold), and low-inflammation ii cluster (blue) (unpaired t-test significant p-value = 0.037). **(E)** Significant taxa binned using tip_glom (phyloseq package in R) identified in pairwise comparisons using a zero-inflated beta regression, plotted over time. Percentage of participants with taxa present in high-inflammation (red) and low-inflammation i (gold) clusters shown in the first three panels (group logarithmic model adjusted p-value ≤ 0.05); abundance (percent of composition) of high-inflammation (red) and low-inflammation ii (blue) clusters shown in fourth panel (group beta regression model adjusted p-value ≤ .05); abundance (percent of composition) of low-inflammation i (gold) and ii (blue) clusters shown in fifth panel (group beta regression model adjusted p-value ≤ 0.05).

Comparison of immune features from baseline to the end of the maintenance phase in the high-fiber diet participants revealed three clusters of participants representing distinct immune response profiles (Figure 4B). These clusters were driven by observed changes in endogenous signaling, most notably decreasing signaling in two clusters (“low-inflammation i” and “low-inflammation ii”), as opposed to an increase in signaling in the “high-inflammation” cluster. Examining individual immune features within the “high-inflammation” cluster revealed increases in JAK/STAT and MAP kinase signaling in monocytes, B cells, and CD4 and CD8 T cells. Both “low-inflammation” clusters showed decreases in these markers (Figure 4C, S4). Taken together, these data suggest divergent immune system responses to the high-fiber intervention, with “high-inflammation” participants exhibiting broad increases in steady-state immune activation versus “low-inflammation” participants exhibiting decreases in steady-state immune activation. Notably, no differences in total fiber intake were observed between inflammation clusters (data not shown).

In order to determine whether these divergent immune system phenotypes were reflected in the participants’ microbiomes, we examined alpha diversity and microbiota composition in the context of the three inflammation clusters. Specifically, a comparison of the observed ASVs at the baseline time points (week -2 and Week 0) for each group revealed higher microbiota diversity in the “low-inflammation ii” group compared to the “high-inflammation” group (p-value=0.037, unpaired t-test) (Figure 4D). While there was not a significant difference between the ‘high-inflammation” and “low-inflammation i” groups (p-value=0.096), there was a trend towards increasing microbiota diversity in the “low-inflammation group i” that followed the intermediate inflammatory response observed. These data are consistent with a previous study demonstrating that a dietary intervention, which included increasing soluble fiber, was less effective in improving inflammation markers in individuals with lower microbiome richness (Cotillard et al., 2013).

A zero-inflated beta regression (ZIBR) model (Chen and Li, 2016) to identify differences in abundance or presence of taxa between clusters over time (Table S7) revealed greater prevalence of *Coprococcus*, *Ruminococcus*, *Oscillospira*, and *Anaerostipes* in the “low-inflammation i” compared to the “high-inflammation” cluster during the high-fiber diet intervention (Figure 4E). *Coprococcus* has been associated with higher quality of life indicators and both *Ruminococcus* and *Oscillospira* have been associated with improved markers of health including leanness and improved lipid profile (Valles-Colomer et al., 2019; Klimenko et al., 2018; Chen et al., 2020). *Anaerostipes* had a significant joint p-value in the ZIBR model (Table S7) and was previously described as a “hyper-butyrate producer” (2014). In contrast, *Akkermansia* was enriched in the “high-inflammation” relative to “low-inflammation ii” cluster. Akkermansia has been positively associated with metabolic health (Derrien et al., 2017), but has also been associated with low-fiber diets and is rare in populations consuming traditional diets (Earle et al., 2015; Desai et al., 2016; Smits et al., 2017). There was a modest difference in a *Lachnospiraceae* taxon between the two low-inflammation clusters. Determining whether a particular taxon is beneficial to human health is problematic given that effects are likely highly context specific and can be influenced by subspecies- (i.e., strain-) specific differences.

### Fermented food intake increases microbiota diversity

In contrast to the high-fiber diet arm, the microbiota of participants consuming the high-fermented foods diet exhibited an overall increase in alpha diversity over the course of the intervention, as determined by overall ASVs and Shannon diversity (Figure 5A, 5B, S2). This diversity increase was sustained during the choice period, when fermented food intake was higher than baseline but lower than at the end of maintenance, suggesting that increased diversity likely involved gut ecosystem remodeling rather than an immediate reflection of consumed quantities. Participants in the high-fermented food diet arm consumed a variety of fermented foods including yogurt, kefir, fermented cottage cheese, kombucha, vegetable brine drinks, and fermented vegetables such as kimchi. Interestingly, while the total number of fermented food servings consumed per day was positively correlated with alpha diversity, the number of servings of yogurt or vegetable brine drinks was most strongly correlated (Figure 5C). Yogurt and vegetable brine drinks were consumed at higher rates relative to the other types of fermented foods, which may contribute to the stronger correlation. Unlike participants consuming the high-fiber diet, no increase in relative abundance of microbial proteins per gram stool was observed in the high-fermented food arm, indicating that altered microbial density did not accompany the increase in diversity (data not shown).

**Figure 5.**
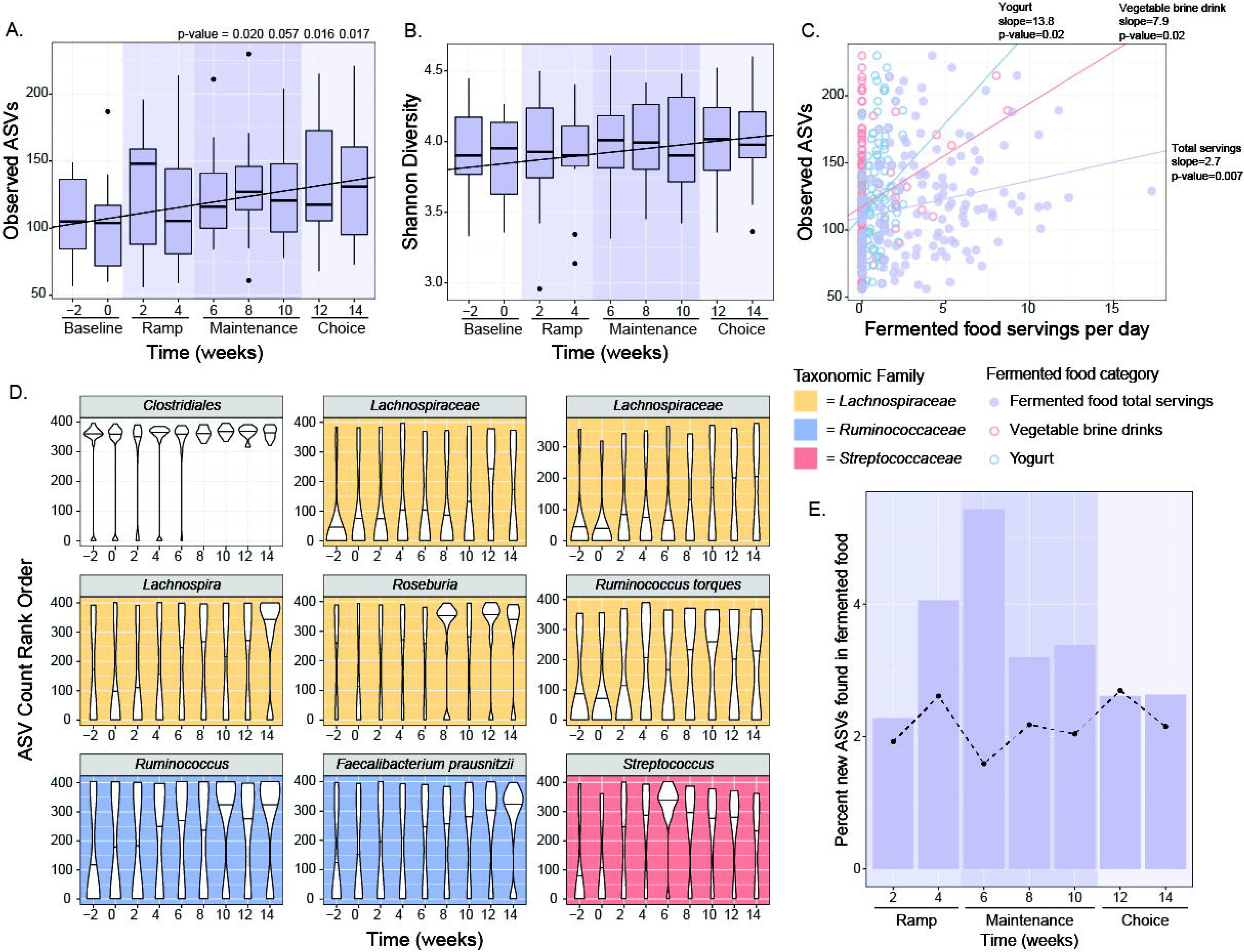
High-fermented food diet increased microbiota diversity and altered composition. **(A, B)** Observed ASVs (A) (p-values generated using paired t-test) and Shannon diversity (B) increased from baseline through choice phase. Observed ASVs significantly correlated with time using linear mixed effects (LME) model (p-value=2.3e-3 for Observed ASVs, p-value=1.4e-3 for Shannon). **(C)** Total fermented food intake, yogurt, and vegetable brine drinks positively correlated with observed ASVs using linear mixed effects (LME) model (p-value adjusted <0.05). **(D)** Rank normalized ASVs that were significantly correlated with fermented food consumption over time using an LME model (p-value adjusted <0.05). Graphs are colored by taxonomic family. **(E)** New ASVs (not present at baseline weeks −2 or 0 but detected at any other time during the intervention) that were detected in fermented foods were aggregated and summed for each participant and plotted as a percentage of all new ASVs by time point for the high-fermented food diet arm. Dotted line indicates trend for high-fiber diet arm.

To examine whether specific taxa changed over time cohort-wide, ASVs were rank-normalized per sample and their relationship with time was modeled using a linear mixed-effect model. Nine ASVs increased over time, all in the Firmicutes phylum, including four in the Lachnospiraceae family, two in the Ruminococcaceae family, and one in the Streptococcaceae family (Figure 5D). An important consideration was whether the new taxa detected were microbes directly sourced from the fermented foods. Taxa present in the commonly consumed fermented foods in the study were identified through 16S rRNA amplicon sequencing (Figure S5) and compared with those newly observed in the participants’ microbiota during the intervention. Only a small percentage of the new microbiota ASVs were common with those found in fermented foods. Peak overlap of new microbiota ASVs and fermented food ASVs occurred early in the intervention, when participants’ overall microbiota diversity was lower than at the end of maintenance phase (5.4%, Fig. 5E). At later time points, overlapping new microbiota ASVs and fermented food ASVs in the fermented food arm was not different than that seen in the high-fiber diet arm (Fig. 5E). These data suggest that the increase in microbiota diversity in the high-fermented food diet arm was not primarily due to consumed microbes, but rather a result of shifts in or new acquisitions to the resident community. These data support that fermented food consumption has an indirect effect on microbiota diversity, rendering the microbiota receptive to the incorporation or increased representation of previously undetected strains within the gut.

The abundance of several CAZyme family members changed from baseline to the end of maintenance phase in the high-fermented food diet arm (Figure S6). However, these changes did not mirror those observed in the high-fiber diet arm. Specifically, all 21 CAZymes that differed in the high-fiber diet arm increased in abundance from baseline to the end of maintenance phase. However, in the fermented food diet arm, the relative abundance of only 10 CAZymes (half of which were annotated as starch-degrading) differed and all decreased in the maintenance phase relative to baseline.

### Fermented food intake decreases markers of host inflammation

Elevated cytokine levels at steady state have been linked to chronic, low-grade inflammation. Using an inflammation panel to assess circulating cytokines in serum, we identified 19 of 93 cytokines, chemokines, and other inflammatory serum proteins that decreased over the fermented food intervention including IL-6, IL-10, and IL-12b and other inflammatory factors (Figure 6A, SAM, FDR < 0.05, q-value < 0.1). IL-6 is a key mediator of chronic inflammation, is elevated in several chronic inflammatory conditions such as rheumatoid arthritis, type-2 diabetes, and chronic stress, and is a commonly used metric of inflammation (reviewed in (Tanaka et al., 2014)). Notably, none of the 19 cytokines that decreased in the high-fermented diet arm were differed in the high-fiber diet arm. We also observed an overall decrease in endogenous signaling, as determined by measuring activation levels of fifteen proteins from four major cell types: CD4+ T cells, CD8+ T cells, B cells, and classical monocytes. Specifically, there were decreased levels of activation in 14 of the 60 different cell type-specific signaling responses, and only one signaling increase (Figure 6B). This decreased signaling was observed across all four cell types tested, consistent with a broad change in immune status in individuals consuming fermented foods. Analysis of CyTOF data to identify the frequency of a larger set of immune cell types revealed that effector memory CD4+ T cells increased, and non-classical monocytes decreased during the intervention (Figure 6C, S2). To evaluate response strength to an immune stimulus, which can be impaired in situations of immune cell exhaustion and aging, we measured the signaling capacity of CD4+ T cells, CD8+ T cells, and B cells in response to *ex vivo* stimulation, but did not find any signaling capacity changes in either diet arm. Together, these data are consistent with an overall cohort-wide decrease in inflammation in the fermented food diet arm.

**Figure 6.**
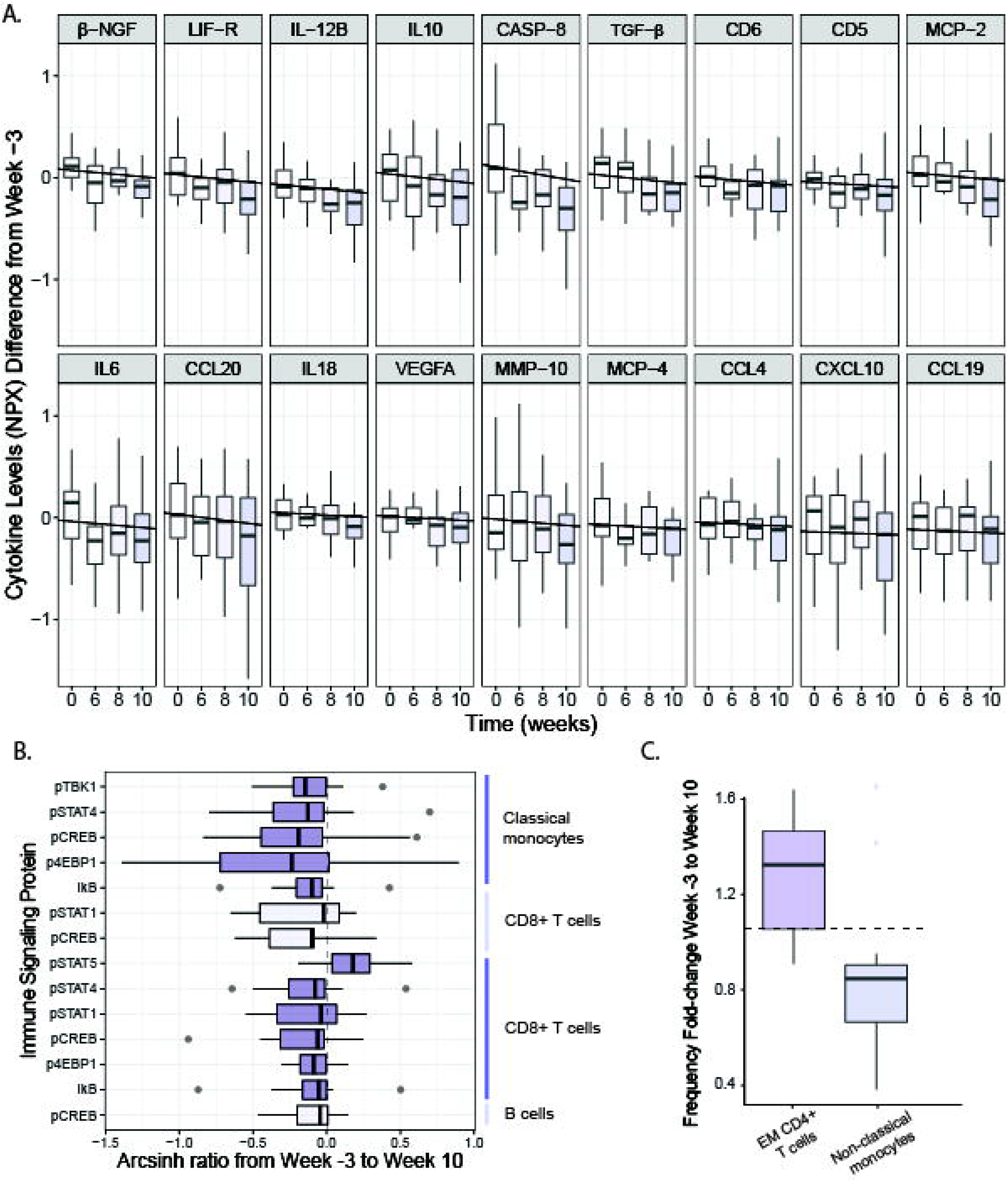
Fermented food consumption decreases levels of inflammation. **(A)** Cytokines, chemokines, and other serum proteins plotted that change significantly from baseline (week -3) to end of intervention (week 10) (SAM two-class paired, FDR ≤ 0.05, q-value ≤ 0.1). Negative correlations for levels of each analyte across time calculated using LME. NPX refers to the normalized protein expression used by Olink Proteomics’ log2 scale. Fgf-21 also significantly decreased across time (data not shown). **(B)** Cell type-specific endogenous signaling proteins, measured using CyTOF, that change significantly from baseline (week -3) to end of intervention (week 10) (SAM two-class paired, FDR < 10%). Arcsinh ratio plotted from week -3 to week 10. **(C)** Fold change of cell frequencies (calculated as percentage of CD45+ cells) that change significantly from baseline (week -3) to end of intervention (week 10) (Wilcoxon paired test, adjusted p-value < 0.05).

### Systems-level microbiome and immune system longitudinal profiling as a method for revealing coordinated host-microbe relationships

Variation in food choices, the individualized nature of participants’ microbiota, and the extensive microbiota and immune system -omics data generated allowed for a unique opportunity to uncover novel human microbiome-immune relationships. To determine the relationship between the microbiota and immune system in a state of change, differences between the end of intervention (Week 10) and baseline (week -3 for blood, week -2 for stool) were calculated for each parameter. These differences were used to determine Spearman correlations between each microbiome feature type (ASVs, alpha diversity, SCFAs, microbe proteomics, stool metabolomics, and CAZymes) and each host feature type (inflammatory cytokines, immune cell signaling, immune cell frequency, and host proteomics). A number of significant correlations (corrected using Benjamini-Hochberg hypothesis correction) between microbiota and host feature type were identified with host stool proteins by microbiota CAZymes having the highest fraction of significant correlations (Figure 7A). Host proteins were annotated and categorized by disease association or function as defined by the Ingenuity Pathway Analysis Core Analysis, Diseases and Functions Analysis (STAR methods, Table S8). The majority of correlations between CAZymes and disease-associated proteins were negative, with proteins assigned to inflammatory response having the highest count of significant correlations (Figure 7B). These associations suggest that CAZymes and host proteins may respond to dietary interventions in coordinated yet opposing directions (i.e. increased CAZyme abundance correlates with decreased levels of inflammation-associated proteins) and may serve as a direct link between diet, the microbiome and host physiology.

**Figure 7.**
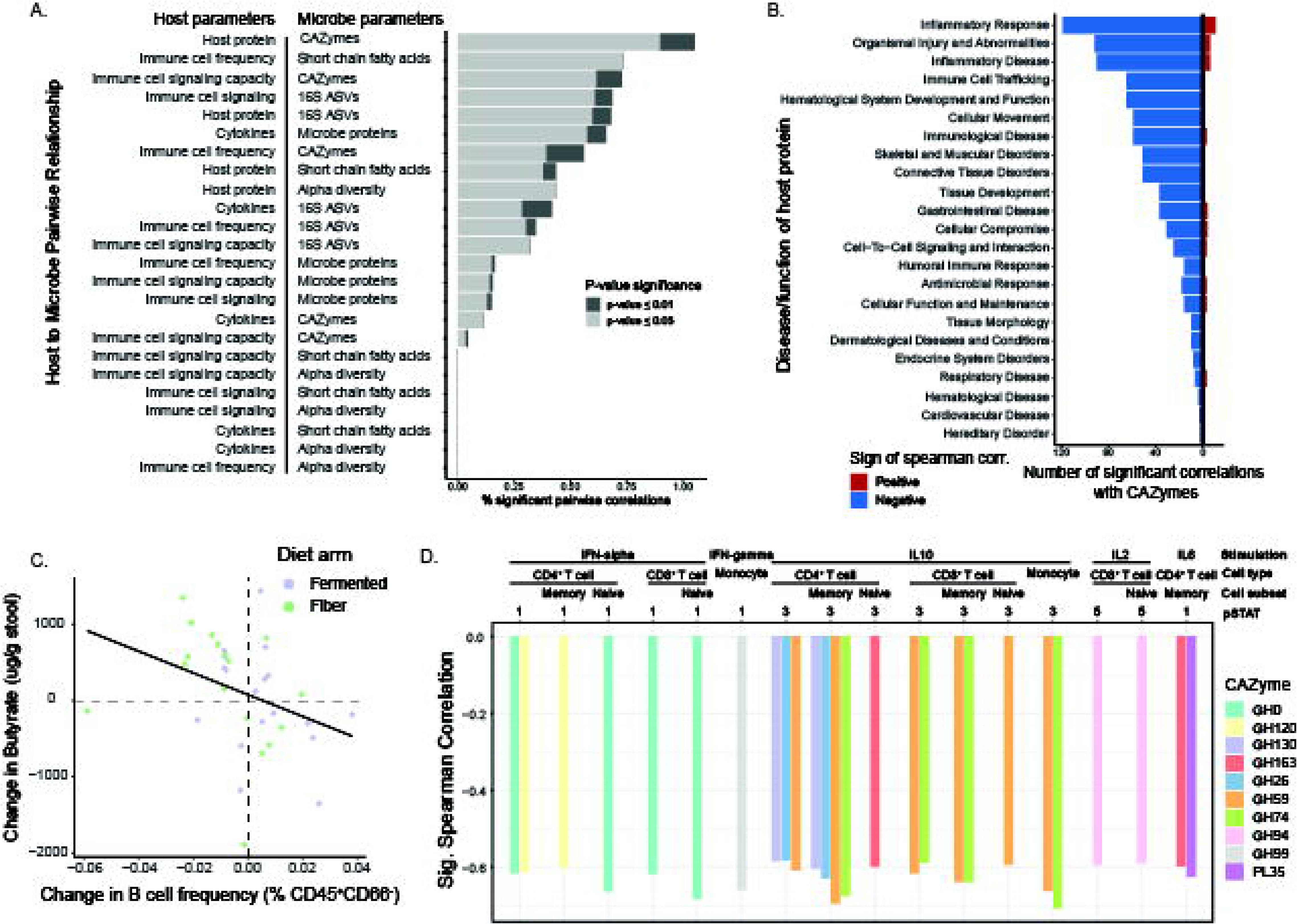
Interaction between the host immune system and microbiota. **(A)** Correlation of difference between baseline and end of maintenance was calculated for each parameter and percent of significant pairwise correlations between microbe and host assays were plotted. Light grey denotes correlations with a p-value adjust < 0.05, dark grey shows p-value adjust < 0.01 (corrected using Benjamini-Hochberg hypothesis correction). **(B)** Positive and negative correlations between host proteins annotated by disease or function (source: Ingenuity Pathway Analysis) and CAZymes. **(C)** Changes in stool butyrate levels vs. blood B-cell frequency changes from baseline (week -3 stool, week -2 blood) to end of maintenance (week 10) for both high-fiber (green) and high-fermented food (purple) arms. **(D)** Correlations between CAZymes (colored by CAZy family) and immune cells signaling capacity for various cell types and stimulatory cytokines.

Our analysis also revealed a relatively high fraction of significant correlations between immune cell frequency and stool short chain fatty acids. Specifically, as fecal butyrate increased, B cell frequency decreased (Figure 7C). B cell depleting therapeutics can be effective treatments for immune-mediated diseases including multiple sclerosis, rheumatoid arthritis, and type 1 diabetes (Fillatreau, 2018). Interestingly, while fecal butyrate did not increase significantly cohort-wide during the high-fiber diet intervention, the majority of participants whose butyrate increased while B cell frequency decreased were in the high-fiber diet arm. Consolidating data from both diet arms revealed this negative association between fecal butyrate and B cell frequency, indicating that a larger cohort may have led to a significant increase in butyrate in the high-fiber diet arm.

The correlations identified between immune cell signaling capacity and microbiome-encoded CAZymes revealed that the abundance of certain CAZymes were negatively correlated with cell signaling capacity (Figure 7D). These data indicate that the apparent attenuation in inflammation indicated by decreased basal level phosphorylation may be further enhanced by CAZyme-linked decreases in cell responsiveness to inflammatory cues. In other words, as participants’ microbiome CAZymes increase in relative abundance they may exhibit a decrease in basal inflammatory status combined with being less responsive after cytokine stimulation.

## Discussion

Extensive data across the field of gut microbiome science has established diet as a major driver of the species and functions that reside within an individual’s gut. Poor diet is a known contributor to non-communicable chronic diseases (NCCDs) that are rapidly spreading globally as more populations adopt Western-style diets (Lozano et al., 2012; GBD 2015 Mortality and Causes of Death Collaborators, 2016). Furthermore, many NCCDs are driven by chronic inflammation, an immunological state that is modulated by the gut microbiota. A logical next phase of gut microbiome science is establishing how diets that influence the gut microbiota modulate human immune status. In this study we used a randomized, prospective dietary intervention model to assess how two components of diet known to interact with the gut microbes, high-fiber and fermented foods, influence the human microbiome and immune system.

Using multiple -omics measurements of the microbiome and host parameters, including state-of-the-art sequencing and immune profiling technologies, we found that high-fiber and high-fermented food consumption influence the microbiome and human biology in distinct ways. One stark difference between these two diets was their impact on gut microbiota diversity. Low microbiota diversity is associated with many NCCDs, such as obesity and diabetes (Turnbaugh et al., 2009; Le Chatelier et al., 2013), and with industrialized lifestyles known to predispose individuals to NCCDs (reviewed in Sonnenburg and Sonnenburg, 2019). Fiber-rich foods contain an abundance of microbiota accessible carbohydrates (MACs), which provide a fermentable carbon source for the microbiota. Despite sustained high levels of diverse plant-derived dietary fiber in these participants over six weeks, we did not observe a cohort-wide microbiota diversity increase in the high-fiber diet arm. It is possible that the relatively short duration of the study was not sufficient to allow for the recruitment of new taxa to the microbiota, which could be an indication that exposure to new microbes was limited within the environment of participants. Environmentally-constrained diversity is consistent with (i) high levels of sanitation in industrialized populations leading to less sharing of microbes between individuals (Martínez et al., 2015), (ii) the necessity of dietary fiber plus administered microbes to restore diversity to the gut microbiota in a mouse model (Sonnenburg et al., 2016), and (iii) the loss of strains and their associated glycan degrading capacity observed in U.S. immigrants (Vangay et al., 2018). The detection of plant glycan-derived carbohydrates in the stool of the high-fiber diet participants is consistent with incomplete microbiota fermentation that might be expected in an industrialized microbiota.

The increased microbiota diversity observed in the fermented food diet arm was coincident with decreases in numerous markers of inflammation, measured with distinct technologies. These correlated changes are consistent with a broad range of studies demonstrating a link between declining microbiota diversity and increased NCCD prevalence (reviewed in Mosca et al., 2016). Notably, the new taxa contributing to the increased diversity were largely not from the fermented foods themselves, indicating an indirect effect of their consumption on remodeling the microbiota. It is unclear whether these “new” taxa were newly recruited to the microbiota from the environment or were already present but undetected and increased in relative abundance to detectable levels during the intervention. The slow trajectory of diversity increase resulted in the greatest microbiota diversity observed during the “choice” phase, where fermented food intake was higher than at baseline, but lower than during the maintenance period. The slow and steadily increasing diversity suggests a time element for remodeling of the microbiome composition through diet, consistent with the relative recalcitrance of the human microbiota to rapid diet-induced remodeling (Wu et al., 2011). Fiber-induced microbiota diversity increases may be a slower process requiring longer than the six weeks of sustained high consumption achieved in this study. Importantly, high-fiber consumption did appear to increase stool microbial protein density, carbohydrate-degrading capacity, and altered SCFA production indicating that microbiome remodeling was occurring within the study time frame, just not through an increase in total species. Given the distinct responses of participants to these two diets, whether a diet composed of both high-fiber and fermented foods could synergize to influence the host microbiota and immune system is an exciting possibility that remains to be determined.

The malleability of the human microbiome, its integration into the immune system, and its responsiveness to diet makes it a highly attractive target for therapeutic intervention. Knowledge of how specific dietary interventions impact the microbiota could be leveraged to develop effective diets that improve human health. Since components of diet, unlike typical pharmaceuticals, do not require regulatory approval for use in humans, they provide an avenue to abate microbiota deterioration and improve human health quickly to avert the coming global NCCD health crisis (GBD 2015 Mortality and Causes of Death Collaborators, 2016). We envision two primary outcomes from additional studies like the ones conducted here. First, precision insight into how one type of dietary intervention may differentially impact individuals, enabling diet to be leveraged in numerous, individual-specific clinical contexts. Second, population-wide insight for diets that broadly improve health, which can serve to guide public health policy, dietary recommendations, and individual choice. For example, fermented food consumption resulted in a cohort-wide generalized dampening in inflammation markers over the course of the intervention. This result is especially striking given that participants in this arm changed little else in their diet and consumed a variety of fermented foods (i.e., some ate mostly fermented dairy products while others ate mostly fermented vegetable products). Additional rigorous studies investigating fermented foods and their impact on human health may lead to the incorporation of these foods as a key component of a healthy diet.

While human studies provide the advantage of illuminating microbiome-host relationships that are relevant to human biology, these come at the cost of mechanistic insight. We view this as a worthwhile trade-off given the ability to reverse translate findings into animal models for mechanistic inquiry based on human-relevant data (Spencer et al., 2019). For example, the host protein and CAZyme relationships we identified here could inform mechanistic mouse studies aimed at understanding the causal relationship between diets that increase carbohydrate utilizing enzymes and decrease inflammatory proteins. In addition, as longitudinal, correlation data (e.g., between microbiome and immune system) for humans accumulates for dietary perturbations, these data can be mined to elucidate a map of interactions between the microbiome and human biology. Such a map will be useful, not only in correcting immune dysregulation that contributes to disease but can be applied in numerous contexts of health and disease to tune biology for optimized physical and mental performance, fighting cancer and numerous chronic diseases, or combatting infectious disease.

## Acknowledgements

We wish to thank the participants for their engagement and effort to enable this study, and Michelle St. Onge for technical support, Biswa P. Choudhury at the UCSD GlycoAnalytics Core for the stool carbohydrates assay, and Susan Kirkpatrick, MS, RD, CDE, Diane Demis, and Erin Avery, MS for assisting with dietary consulting. We would also like to thank Kavita Mathi and Natalia Sigal from the Human Immune Monitoring Core for staining and running phosphoflow and CyTOF assays. This work was funded by generous donations to the Center for Human Microbiome Research, Paul and Kathy Klingenstein, the Hand Foundation, Heather Buhr and Jon Feiber, and Meredith and John Pasquesi. G.K.F. was supported by NIH T32 AI 7328-29 and the Stanford Dean’s Postdoctoral Fellowship. H.C.W. was supported by the NSF Graduate Student Fellowship. J.L.S is a Chan-Zuckerberg Biohub investigator.

## Author contributions

H.C.W, G.K.F., M.T., J.L.S., C.D.G., and E.D.S. performed the experiments and designed and performed data analyses. D.P. provided patient interaction, dietary counseling and analysis. D.D., B.D.M., and F.B.Y. led metagenomics experiments and analysis. C.G.G. and J.E.E. designed and performed stool proteomics experiments and guided their analysis. J.L.R. oversaw patient recruitment and study design implementation. H.C.W., G.K.F., J.L.S., C.D.G. and E.D.S. conceived of the study and wrote the manuscript.

## Supplemental figure titles and legends

**Figure S1, Related to Figure 1:** Individual participant high-fiber and high-fermented food diet arm group intake.

**(A)** Participant-specific fiber intake by category for high-fiber diet arm.

**(B)** Participant-specific intake by category for high-fermented foods diet arm.

**Figure S2, Related to Figures 4 and 6:** CyTOF gating strategy

**Figure S3, Related to Figures 3 and 5:** Alpha diversity measures for high-fiber and high-fermented diet arms.

**(A)** Shannon alpha diversity in high-fiber diet arm.

**(B)** Phylogenetic diversity (PD) whole tree alpha diversity in high-fiber food diet arm.

**(C)** PD whole tree alpha diversity in high-fermented food diet arm. * indicates significant p-value relative to week -2 (p-value < 0.05). Linear regression significant using a linear mixed effects model (p-value=0.013)

**Figure S4, Related to Figure 4:** Changes in endogenous signaling in the high-fiber diet arm inflammation groups. Endogenous signaling levels measured by CyTOF and identified as significantly changed (FDR ≤ 0.05, q-value ≤ 0.1, SAM test using siggenes package) from baseline (week -3) to end of maintenance (week 10) indicated by the shaded boxes next to the boxplots (red=significantly changed in high-inflammation group, blue=significantly changed in low-inflammation ii group).

**Figure S5, Related to Figure 5:** 16S ASV analysis of fermented foods. ASVs with fewer than 250 counts were filtered out and counts binning to the same genus were summed. If genus was unassigned, ASVs were summed based on the same order and family.

**Figure S6, Related to Figure 5:** CAZyme analysis of high-fermented food diet arm. CAZymes identified from metagenomic sequencing as significantly changing in relative abundance from baseline to end of maintenance phase (FDF < 0.05, q-value < 0.1, SAM test using siggenes package). CAZymes were annotated using dbCan and assigned to functional categories.

## Supplemental table titles and legends

**Table S1, Related to Figure 1:** Demographics table

**Table S2, Related to Figure 1:** Primary and secondary clinical trial outcomes

**Table S3, Related to Figure 1:** Number of participants per data type

**Table S4, Related to Figure 1:** Nutrient data from participant diets. A) High-fiber diet arm nutrient intake. B) High-fermented diet arm nutrient intake

**Table S5, Related to Figure 2:** Features selected in random forest models predicting diet group

**Table S6, Related to Figure 4 and 6:** Immune profiling panels. Contains panels for Olink, CyTOF surface makers, CyTOF intracellular markers, and Phospho-flow

**Table S7, Related to Figure 4:** ZIBR coefficients for high-fiber diet arm inflammation clusters. ZIBR coefficients on the tip glommeration for fiber inflammation groups. Beta-regression model refers to the change in relative abundance of taxa. Logarithmic model refers to presence/absence of taxa. A significant coefficient for the grouping variable indicates a significant difference between the inflammation groups indicated in the comparison column. Baseline coefficients indicate a difference in the taxonomic relative abundance or presence/absence at the baseline time point (week 0). Taxa with significant baseline coefficients were filtered out to focus on the significant differences induced by the dietary intervention.

**Table S8, Related to Figure 7:** Protein disease annotations.

## Materials and Methods

### RESOURCE AVAILABILITY

#### Data and Code Availability

Datasets and code for analysis are available at https://github.com/SonnenburgLab/fiber-fermented-study/.

### EXPERIMENTAL MODEL AND SUBJECT DETAILS

#### Recruitment and selection of participants

Participants were recruited from the local community through online advertisement in different community groups as well as emails to past research participants that consented to being contacted for future studies. The current study assessed 381 participants for eligibility. They completed an online screening questionnaire and a clinic visit between July 2016 and January 2017. The primary inclusion criteria included age ≥ 18 y and general good health. Participants were excluded if they had a history of active uncontrolled inflammatory bowel disease (IBD) including ulcerative colitis, Crohn’s disease, or indeterminate colitis, irritable bowel syndrome (IBS) (moderate or severe), infectious gastroenteritis, colitis or gastritis, *Clostridium difficile* infection (recurrent) or *Helicobacter pylori* infection (untreated), malabsorption (such as celiac disease), major surgery of the GI tract, with the exception of cholecystectomy and appendectomy, in the past five years, or any major bowel resection at any time. Other exclusion criteria included a BMI ≥ 40, diabetes, renal disease, significant liver enzyme abnormality, pregnancy or lactation, smoking, a history of CVD, inflammatory disease, or malignant neoplasm. Participants with high levels of dietary fiber intake (above 20 g of fiber per day) or more than 2 servings per day of fermented foods were excluded. Consort flow diagram of participant recruitment shown in Figure 1A and demographics table shown in Table S1. 36 participants (25 female sex and gender identifying, 11 male sex and gender identifying) were used for full analysis with an average age of 52 +/− 11 years. All study participants provided written informed consent. The study was approved annually by the Stanford University Human Subjects Committee. Trial was registered at ClinicalTrials.gov, identifier: NCT03275662

#### Specimen collection

Stool samples were collected every two weeks from week -2 through the end of observation at week 14. All stool samples were kept in participants’ home freezers (−20°C) wrapped in ice packs, until they were transferred on ice to the research laboratory and stored at −80°C.

Blood samples were collected at 7 time points: −3 weeks, start of intervention, week 4 (end of ramp-up), week 6, week 8, week 10 (3 time points during maximum intake), and week 14 (end of observation). Blood for PBMC and whole blood aliquots were collected into heparinized tubes. Whole blood aliquots were incubated with Proteomic Stabilization Buffer (Smart tube, Fisher Scientific) for 12 minutes at room temperature and stored at −80C. PBMCs were isolated using Ficoll-Paque PLUS (Sigma-Aldrich), washed with PBS, frozen at −80C for 24 hours then moved to LN2 for longer storage. Blood for serum was collected into an SST-tiger top tube, spun at 1,200xg for 10 minutes, aliquoted, and stored at −80C. Blood for plasma was collected into an EDTA tube, spun at 1,200xg for 10 minutes, aliquoted, and stored at −80C.

### METHOD DETAILS

#### Intervention

Participants were randomized to follow a diet high in fiber or high in fermented foods. They were instructed to ramp up the intake of foods high in fiber/fermented during the first 4 weeks of the intervention with a goal of increasing 20 g/day of fiber in the fiber arm and 6 servings a day of fermented foods/day in the fermented food arm, and were encouraged to consume more if they could tolerate it. They were instructed to maintain the high level of consumption during the following 6 weeks. Detailed instructions were provided to encourage participants to include a variety of fiber sources (legumes, seeds, whole grains, nuts, vegetables, and fruits) or fermented foods (fermented dairy products, fermented vegetables, fermented non-alcoholic drinks). Participants were followed for an additional 4 weeks after the end of the intervention period. All participants met with a dietitian at baseline, end of ramp up, and every 2 weeks during the high intake period. They were asked to keep detailed food logs 3 days per week (2 weekdays and 1 weekend) every other week through the duration of the study. Food logs were reviewed by the dietitian to assess compliance and provide recommendations to increase amounts or variety of fiber/fermented foods in the diet as tolerated. Participants filled out gastrointestinal symptoms surveys (GSRS) (Svedlund et al., 1988) and symptom changes (Winham and Hutchins, 2011) every 2 weeks, and these were discussed during the visits with the dietitian.

#### Dietary Data

Participants logged all their food and drink intake for 3 days (2 weekdays and 1 weekend) each week during the ramp phase and every other week for the rest of the study using the HealthWatch360 app. The dietitian reviewed the entries with participants to assess accuracy of entries and portions. An average of the 3 days was used for each time point. Entries for time points: baseline, week 4, week 10, and week 14 were re-entered into the Nutrition Data System for Research (NDSR) 2019 database by a dietitian. NDSR appendix 10 was used to classify foods into food groups.

Fiber intake was reported as average intake in grams per day for each week of the intervention. Categories of fiber sources were grouped into fruits, grains, legumes, nuts/seeds, vegetables, meat, dairy, and other. Fermented intake was reported as the average number of servings per day for each week of the intervention. One serving of fermented foods were defined as the following: kombucha, yogurt, kefir, buttermilk, kvass = 6 oz, kimchi, sauerkraut, other fermented veggies = 1/4 cup, vegetable brine drink = 2 oz. To determine if fiber or fermented intake significantly changed during the course of the intervention, paired t-tests were performed from week -2 to all other time points. Broad categories of fermented foods were grouped into cottage cheese, kefir, kombucha, vegetable brine drinks, vegetables, yogurt, other foods, and other drinks.

The following validated health surveys were used by participants: PROMIS v1.1 global health, PROMIS v1.0 - fatigue, WHO well-being index, PROMIS applied cognition short form, Perceived Stress Scale (Cohen et al., 1983), and the International Physical Activity Questionnaire (Craig et al., 2003).

#### 16S amplicon sequencing

DNA was extracted from stool and fermented foods using the MoBio PowerSoil kit according to the Earth Microbiome Project’s protocol (Gilbert et al., 2014) and amplified at the V4 region of the 16S ribosomal RNA (rRNA) subunit gene and 250 nucleotides (nt) Illumina sequencing reads were generated. There was an average of 20,119 reads per sample and samples with less than 1,000 reads were filtered out (7 samples out of 338 removed). There was an average of 15,292 reads per sample recovered after filtering, denoising, and removing chimeras. Fermented foods of the same brands used commonly by participants were purchased and subjected to the 16S sequencing method. It is important to note that our findings for these foods may not directly reflect the composition of the exact foods (e.g., due to batch variation) eaten by the participants, who purchased their own fermented food.

16S rRNA gene amplicon sequencing data from both stool samples and fermented food samples were demultiplexed using the QIIME pipeline version 1.8 (Caporaso et al., 2010). Amplicon sequence variants (ASVs) were identified with a learned sequencing error correction model (DADA2 method) (Callahan et al., 2016), using the dada2 package in R. ASVs were assigned taxonomy using the GreenGenes database (version 13.8).

α-diversity was quantified as the number of observed ASVs, Shannon diversity, or PD whole tree, in a rarefied sample using the phyloseq package in R (version 3.4.0). Data were rarefied to 3,649 reads per sample (lowest 10% of reads, 299 samples retained out of 331 total) also using the phyloseq package in R. Rarefied data were only used for α-diversity measures.

#### Metagenomic sequencing

DNA extraction for shotgun metagenome sequencing was done using the MoBio PowerSoil kit as described in the 16S amplicon sequencing methods. For library preparation, the Nextera Flex kit was used with a minimum of 10ng of DNA as input and 6 or 8 PCR cycles depending on input concentration. A 12 base pair dual-indexed barcode (CZ Biohub) was added to each sample and libraries were quantified using an Agilent Fragment Analyzer. They were further size-selected using AMPure XP beads (Beckman) targeted at a fragment length of 450bp (350bp size insert). DNA paired-end sequencing (2×146bp) was performed on a NovaSeq 6000 using S4 flow cells (CZ Biohub). The average target depth for each sample was 23.3 million paired-end reads.

Data quality analysis was performed by demultiplexing raw sequencing reads and concatenating data for samples that required multiple sequencing runs for target depth before further analysis. BBtools suite (https://sourceforge.net/projects/bbmap/)) was used to process raw reads and mapped against the human genome (hg19) after trimming, with masks over regions broadly conserved in eukaryotes (http://seqanswers.com/forums/showthread.php?t=42552). Exact duplicate reads (subs=0) were marked using clumpify and adapters and low-quality bases were trimmed using bbduk (trimq=16, minlen=55). Finally, reads were processed for sufficient quality using FastQC (https://www.bioinformatics.babraham.ac.uk/projects/fastqc/).

Carbohydrate active enzymes (CAZymes) were annotated using dbCAN (v2.0.11) (Yin et al., 2012; Huang et al., 2018) on genes called from FragGeneScan (Rho et al., 2010). From merged reads, unmerged reads with the requirement that the CAZymes were identified with both diamond (Buchfink et al., 2015) and hotpep (Busk et al., 2017). Final read counts were normalized by calculating the reads per million for each CAZyme subfamily in each of the samples (CAZyme count/sum all sample counts/1e6). CAZyme analysis was restricted to GHs and PLs. To determine the CAZyme subfamilies that significantly changed in relative abundance from baseline to end of maintenance phase, a using the siggenes package in R (SAM two-class paired, FDR ≤ 0.05, q-val ≤ 0.01).

#### Stool proteomics

Methods for stool preparation, mass spectrometry protocol, and protein searches are described in Gonzalez et. al. Briefly, a measured quantity between 100-200 mg of stool per sample was loaded into a 96-well plate and lysed with ceramic beads, centrifuged, supernatant proteins alkylated, washed, digested and eluted using an S-trap plate. Protein concentrations were normalized and longitudinal samples for each individual labeled with TMT-11 multiplexing kit. Peptide samples were injected onto reversed-phase chromatography using a Dionex Ultimate 3000 HPLC and run on a Thermo Fusion Lumos mass spectrometer that collected MS data in positive ion mode within a 400-1500 m/z range. The resulting mass spectra raw data files were searched using Proteome Discoverer 2.2. using the built-in SEQUEST search algorithm with built-in TMT batch correction. Three FASTA protein sequence databases were employed: Uniprot Swiss-Prot *Homo sapiens* (taxon ID 9606, FASTA file downloaded January 2017), the Human Microbiome Project (FASTA file downloaded from https://www.hmpdacc.org/hmp/HMRGD/ on January 2017), and an in-house curated database containing common preparatory contaminants. Target-decoy searching at both the peptide and protein level were employed with a strict FDR cutoff of 0.01 using the Percolator algorithm built into Proteome Discoverer 2.2. The mass spectrometry proteomics data have been deposited to the ProteomeXchange Consortium via the PRIDE partner repository with the data set identifier PXD021786.

Protein abundance was recorded from the average of two runs, log2(x+1) transformed, and normalized as a percentage of summed reporter intensity for all quantified proteins in a given sample (single protein intensity/total sample intensity). Proteins were denoted as either host or microbial derived based on high confidence matches to the Human Microbiome Project protein sequence database. Reported microbial proteins and protein abundances represent the summation of protein database entries with identical descriptions, decreasing microbial protein variables from 5,372 unique proteins to 4,315 unique descriptions. All host proteins had unique descriptions. As an unsupervised method to decrease the number of parameters for multiple hypothesis testing, the host proteins were filtered to include the top 75% (230 proteins) and microbe proteins were filtered to include the top 50% (2,157 proteins) proteins with the highest variance across participants (participant-specific difference from end of intervention to baseline). All proteomic analyses were completed using the described filtered data set.

#### Stool short-chain fatty acids

For sample preparation, a measured quantity of ~20 mg stool per sample was loaded into 1.5 mL eppendorf tubes and sent to Metabolon for absolute quantitation of short chain fatty acids. All samples were kept frozen and shipped on dry ice. Paired Welsh t-tests were used to determine if levels of short chain fatty acids significantly changed from baseline to end of intervention.

#### Stool Carbohydrates

Methods for measurement of monosaccharides present in stool samples using GC-MS are described in (DeJongh et al., 1969). Briefly, stool samples were homogenized in 10% aqueous ethanol solution and dried using a vacuum concentrator. A known dry weight (between 0.5-0.8mg) of sample was transferred to a glass hydrolyzing tube before being suspended via sonication, heated, and lyophilized. Samples were then methanolyzed using 1M MeOH-HCl and re-N-acetylated using MeOH: Pyridine: Acetic anhydride (3:1:1 v/v). Finally, samples were converted to TMS-ehteress using Tri-Sil reagent (Thermo Scientific), dried via dry nitrogen flush, and extracted with hexane. TMS derivatives of the methyl glycosides were analyzed using GC-MS and profiling of the monosaccharides was completed using Resteck 5MS fused silica capillary column at an oven temperature gradient. 1uL of sample was injected onto the GC column using split-less mode. A standard mixture of different monosaccharides was also run to compare and quantify absolute value of monosaccharides present in samples. A linear mixed effects model was used to vary percentage of carbohydrates in stool with fiber intake.

#### CyTOF

Whole blood samples were thawed and red blood cells were lysed using Thaw Lyse buffer (Smart Tube, inc) for 10 minutes twice at room temperature, then washed twice with cell staining media (CSM: PBS with 0.5% BSA and 0.02% Na azide). 1×10^6^ cells from each sample were barcoded as previously described (Behbehani et al., 2014). Briefly, cells were slightly permeabilized using PBS with 0.02% saponin, then stained with unique combinations of functionalized, stable palladium isotopes for 15 minutes at room temperature. Samples were washed with CSM and pooled into a single tube for staining. Cells were blocked with human TruStain FcX block (Biolegend) then stained with an extracellular antibody cocktail. Antibody cocktails were rehydrated with CSM after lyophilization into LyoSpheres (BioLyph) with excipient B144 as 4x cocktails as previously described (Fragiadakis et al., 2019). Antibody panels are listed in Table S6. Samples were then permeabilized using methanol for 10 minutes at 4°C and stained with an intracellular antibody cocktail. Cells were stained with an iridium intercalator overnight prior to CyTOF acquisition. Samples were washed twice with water, resuspended in normalization beads (Fluidigm), and filtered through a cell strainer. Samples were run on a Helios CyTOF.

Samples were normalized and debarcoded using the premessa package in R. Cell populations were gated using Cell Engine (immuneatlas.org, gating strategy, Figure S2). Cell frequencies were calculated as the fraction of CD45+ cells, with the exception of neutrophils, which were quantified as a percentage of singlet cells. Endogenous signaling was taken as the median level of the transformed level of a signaling protein in a specific cell population (transformation = arcsinh(value/5)). Samples were excluded if the number of singlets was less than 10,000 cells. If data were available for both baseline samples, the first baseline was used; otherwise the second baseline was used. For heatmaps, missing data were imputed using the average value of a feature across all participants. For significance analysis, participants with missing data were excluded. For significance analysis of signaling proteins, features were restricted to those in four major cell types (CD4+ T cells, CD8+ T cells, B cells, classical monocytes) and analysis was performed using the siggenes package in R (SAM two-class paired, FDR ≤ 0.05, q-value ≤ 0.1). Significance of cell frequencies was assessed using a Wilcoxon paired test.

#### Serum cytokines

Cytokine data were generated from serum samples submitted to Olink Proteomics for analysis using their provided inflammation panel assay of 92 analytes (Olink INFLAMMATION,Table S6). Out of 92 proteins, 67 were detected in >75% of samples and used in analysis. Data are presented as normalized protein expression values (NPX, Olink Proteomics arbitrary unit on log2 scale). Significance was assessed using the siggenes package in R (SAM two-class paired, FDR ≤ 0.05, q-value ≤ 0.1).

#### Flow cytometry

This assay was performed by the Human Immune Monitoring Center at Stanford University. PBMC were thawed in warm media, washed twice and resuspended at 0.5×10^6^ viable cells/mL. 200 uL of cells were plated per well in 96-well deep-well plates. After resting for 1 hour at 37°C, cells were stimulated by adding 50 ul of cytokine (IFNa, IFNg, IL-6, IL-10, or IL-2) or LPS and incubated at 37°C for 15 minutes. The PBMCs were then fixed with paraformaldehyde, permeabilized with methanol, and kept at −80°C overnight. Each well was bar-coded using a combination of Pacific Orange and Alexa-750 dyes (Invitrogen, Carlsbad, CA) and pooled in tubes. The cells were washed with FACS buffer (PBS supplemented with 2% FBS and 0.1% sodium azide), and stained with the following antibodies (all from BD Biosciences, San Jose, CA): CD3 Pacific Blue, CD4 PerCP-Cy5.5, CD20 PerCp-Cy5.5, CD33 PE-Cy7, CD45RA Qdot 605; cytokine samples were additionally stained with pSTAT-1 FITC, pSTAT-3 APC, pSTAT-5 PE, whereas the LPS sample was stained with pERK APC, pP38 FITC, and pPLCg2 PE (Table S6). The samples were then washed and resuspended in FACS buffer. 100,000 cells per stimulation condition were collected using DIVA 6.0 software on an LSRII flow cytometer (BD Biosciences). Gating was performed using FlowJo v9.3 by gating on live cells based on forward versus side scatter profiles, then on singlets using forward scatter area versus height, followed by cell subset-specific gating.

Signaling markers were quantified as the 90th percentile value. To quantify signaling capacity, we calculated fold change in phospho-proteins between cytokine stimulated and unstimulated. For analysis we restricted our feature set to features with an average fold change greater than two. Significance was assessed using the siggenes package in R (SAM two-class paired, FDR ≤ 0.05, q-value ≤ 0.1).

### QUANTIFICATION AND STATISTICAL ANALYSIS

#### Location of Statistical Details in the Text

Results of each experiment can be found in the results and figure legends. Significant values of statistical tests are also indicated by an asterisk in the figures. Number of participants in each data type are found in Supplementary Table 1.

#### Centering and Scaling Data

In analyses comparing different data types to each other, parameters were centered and scaled (across columns) to eliminate experimental bias. All methods describing data as centered and scaled were done using the scale function in base R (scale function, center=TRUE, scale=TRUE). Centered data were calculated by subtracting the column means from each value. Scaled data were calculated by dividing the centered columns by their standard deviations.

#### Recursive feature random forest

To determine which data type was most accurate in differentiating between high-fiber and high-fermented food diet arms, a recursive feature random forest (caret, rfeControl, number=100, leave one out cross validation) was used. Parameters for each data type were the participant-specific differences from end of intervention (week 10) to baseline (week -2 for stool, week -3 for blood). If a participant did not have both the baseline and end of intervention time point for a given experimental platform they were removed from analysis. All parameters were centered and scaled. To decrease redundant parameters of large feature sets, unsupervised parameter filtration was used. 16S data were filtered to only ASVs present in at least 25% of samples and rank-normalized within each sample according to the methods described by (“Workflow for Microbiome Data Analysis: from raw reads to community analyses.,” n.d.). Host proteins were filtered to top 75% (230 proteins total) and microbe proteins were filtered to top 50% proteins (2,157 proteins total) with the highest variance across participants. CAZyme subfamilies were summed at the family level across samples. The recursive feature random forest models returned the minimum feature set needed for highest accuracy (Table S5).

#### Multiple Testing using Significance Analysis of Microarrays (SAM)

The identification of parameters differentially expressed between diet groups (unpaired) or within the same participant at different time points (paired) and estimation of the False Discovery Rate (FDR) was calculated using the siggenes package in R. Significance was described as FDR ≤ 0.05 and a q-value ≤ 0.10.

#### Linear Mixed Effects Modeling

Linear mixed-effects models were used to assess the linear correlation between two variables when the same participant contributed multiple samples to the model (i.e., Participant at multiple time points). Because samples from the same participant are not independent from one another and introduces autocorrelation, we used the participant term as a random variable in the lme function using the nLME package in R.

Total fiber intake (grams) was correlated with the percentage of carbohydrates in stool using a linear mixed-effects model using the lme function and participants as the random variable.

Association between rank-order ASV count vs. time point (weeks) and alpha diversity (number observed ASVs) vs. fermented food intake were assessed using the lme function and participants as the random variable. P-values for all ASVs in association with time were adjusted for multiple hypothesis testing using a Benjamini-Hochberg correction.

To determine if the number of observed ASVs over time varied between high-fiber diet inflammation groups, a pairwise LME model was used. Number of observed ASVs was the outcome variable with both time (in weeks) and inflammation group (binary variable) as the covariates. The inflammation group only compared two groups at a time, so three models in total were made to determine significance between fiber inflammatory groups: high-inflammation vs. low-inflammation i, high-inflammation vs. low-inflammation ii, and low-inflammation i vs. low-inflammation ii. For the model comparing high-inflammation and low-inflammation i, the inflammation group factor was significant (p-value = 7.2e-4), but the time variable was not (p-value = 0.69). The other two models did not have significant p-values.

To determine if fermented food intake varied significantly with the number of observed ASVs, the lme function and participants as the random variable, a model was made for total fermented food intake and each fermented food group separately, p-values were adjusted using Benjamini-Hochberg correction.

#### Modeling ASV Changes in Relative Abundance and Presence/Absence Over Time using a Zero-inflated Beta Random Effect Model (ZIBR)

To identify differences in abundance and/or presence of taxa between inflammation clusters over time in the high-fiber diet arm, the zero-inflated beta regression model was fit using the ZIBR package in R. A filtered dataset was curated as described in (Chen and Li, 2016). ASVs were preprocessed using tip_glom (phyloseq package in R, h=0.1), removed if they were non-characterized in GreenGenes, and filtered to only ASVs present in at least 25% of samples. Since ZIBR cannot handle missing data, missing samples were filled with the average ASV abundance for each group at each timepoint. Taxa with significant baseline coefficients were filtered out to focus on the significant differences induced by the dietary intervention (Table S7).

#### New ASVs in Participant Samples also Detected in Fermented Foods

New ASVs in participant stool samples were calculated by aggregating and summing new ASVs for each participant and time point. New ASVs included those not present at either baseline time point (weeks -2, 0), but detected at any other time point during the intervention (weeks 2-9). Fermented food ASVs (Table S5) with less than 250 counts were filtered out. Overlap of the new ASVs gained during intervention and also found in the fermented food were summed across all participants at each time point.

#### Spearman Correlation Between Data Types

The spearman correlation between all parameters was calculated. Data input encompassed participant-specific differences from both groups, centered and scaled. If a participant did not have both the baseline and end of intervention time point for a given experimental platform they were removed from analysis. Parameters were filtered using the same methods described for the random forest, grouped into their respective experimental platforms, and designated to host or immune derived. Correlations were filtered to only host-microbe comparisons before Benjamini-Hochberg hypothesis correction. Correlations between host proteins and microbe proteins are not shown here as they were derived from the same sample and experimental platform and had inflated internal bias compared to the other cross-omic comparisons. Host protein annotation for analysis with their association to CAZymes (Figure 7B) was assigned using the Ingenuity Pathway Analysis Core Analysis, Diseases and Functions Analysis. Full annotation of proteins in each bin can be found in Table S8.

### ADDITIONAL RESOURCES

Clinical trial registry #NCT03275662: https://clinicaltrials.gov/ct2/show/NCT03275662

## References

Arumugam, M., Raes, J., Pelletier, E., Le Paslier, D., Yamada, T., Mende, D.R., Fernandes, G.R., Tap, J., Bruls, T., Batto, J.-M., et al. (2011). Enterotypes of the human gut microbiome. Nature 473, 174–180.

Behbehani, G.K., Thom, C., Zunder, E.R., Finck, R., Gaudilliere, B., Fragiadakis, G.K., Fantl, W.J., and Nolan, G.P. (2014). Transient partial permeabilization with saponin enables cellular barcoding prior to surface marker staining. Cytom. Part J. Int. Soc. Anal. Cytol. 85, 1011–1019.

Benesh, A.E., Nambiar, R., McConnell, R.E., Mao, S., Tabb, D.L., and Tyska, M.J. (2010). Differential localization and dynamics of class I myosins in the enterocyte microvillus. Mol. Biol. Cell 21, 970–978.

Brand-Miller, J., and Buyken, A. (2020). Mapping postprandial responses sets the scene for targeted dietary advice. Nat. Med. 26, 828–830.

Buchfink, B., Xie, C., and Huson, D.H. (2015). Fast and sensitive protein alignment using DIAMOND. Nat. Methods 12, 59–60.

Busk, P.K., Pilgaard, B., Lezyk, M.J., Meyer, A.S., and Lange, L. (2017). Homology to peptide pattern for annotation of carbohydrate-active enzymes and prediction of function. BMC Bioinformatics 18, 214.

Callahan, B.J., McMurdie, P.J., Rosen, M.J., Han, A.W., Johnson, A.J.A., and Holmes, S.P. (2016). DADA2: High-resolution sample inference from Illumina amplicon data. Nat. Methods 13, 581–583.

Cantarel, B.L., Lombard, V., and Henrissat, B. (2012). Complex carbohydrate utilization by the healthy human microbiome. PloS One 7, e28742.

Caporaso, J.G., Kuczynski, J., Stombaugh, J., Bittinger, K., Bushman, F.D., Costello, E.K., Fierer, N., Peña, A.G., Goodrich, J.K., Gordon, J.I., et al. (2010). QIIME allows analysis of high-throughput community sequencing data. Nat. Methods 7, 335–336.

Chen, E.Z., and Li, H. (2016). A two-part mixed-effects model for analyzing longitudinal microbiome compositional data. Bioinforma. Oxf. Engl. 32, 2611–2617.

Chen, Y.-R., Zheng, H.-M., Zhang, G.-X., Chen, F.-L., Chen, L.-D., and Yang, Z.-C. (2020). High Oscillospira abundance indicates constipation and low BMI in the Guangdong Gut Microbiome Project. Sci. Rep. 10, 9364.

Cohen, S., Kamarck, T., and Mermelstein, R. (1983). A global measure of perceived stress. J. Health Soc. Behav. 24, 385–396.

Cotillard, A., Kennedy, S.P., Kong, L.C., Prifti, E., Pons, N., Le Chatelier, E., Almeida, M., Quinquis, B., Levenez, F., Galleron, N., et al. (2013). Dietary intervention impact on gut microbial gene richness. Nature 500, 585–588.

Craig, C.L., Marshall, A.L., Sjöström, M., Bauman, A.E., Booth, M.L., Ainsworth, B.E., Pratt, M., Ekelund, U., Yngve, A., Sallis, J.F., et al. (2003). International physical activity questionnaire: 12-country reliability and validity. Med. Sci. Sports Exerc. 35, 1381–1395.

David, L.A., Maurice, C.F., Carmody, R.N., Gootenberg, D.B., Button, J.E., Wolfe, B.E., Ling, A.V., Devlin, A.S., Varma, Y., Fischbach, M.A., et al. (2014). Diet rapidly and reproducibly alters the human gut microbiome. Nature 505, 559–563.

Deehan, E.C., and Walter, J. (2016). The Fiber Gap and the Disappearing Gut Microbiome: Implications for Human Nutrition. Trends Endocrinol. Metab. TEM 27, 239–242.

De Filippo, C., Cavalieri, D., Di Paola, M., Ramazzotti, M., Poullet, J.B., Massart, S., Collini, S., Pieraccini, G., and Lionetti, P. (2010). Impact of diet in shaping gut microbiota revealed by a comparative study in children from Europe and rural Africa. Proc. Natl. Acad. Sci. U. S. A. 107, 14691–14696.

DeJongh, D.C., Radford, T., Hribar, J.D., Hanessian, S., Bieber, M., Dawson, G., and Sweeley, C.C. (1969). Analysis of trimethylsilyl derivatives of carbohydrates by gas chromatography and mass spectrometry. J. Am. Chem. Soc. 91, 1728–1740.

Derrien, M., Belzer, C., and de Vos, W.M. (2017). Akkermansia muciniphila and its role in regulating host functions. Microb. Pathog. 106, 171–181.

Desai, M.S., Seekatz, A.M., Koropatkin, N.M., Kamada, N., Hickey, C.A., Wolter, M., Pudlo, N.A., Kitamoto, S., Terrapon, N., Muller, A., et al. (2016). A Dietary Fiber-Deprived Gut Microbiota Degrades the Colonic Mucus Barrier and Enhances Pathogen Susceptibility. Cell 167, 1339–1353.e21.

Díaz-López, A., Bulló, M., Martínez-González, M.A., Corella, D., Estruch, R., Fitó, M., Gómez-Gracia, E., Fiol, M., García de la Corte, F.J., Ros, E., et al. (2016). Dairy product consumption and risk of type 2 diabetes in an elderly Spanish Mediterranean population at high cardiovascular risk. Eur. J. Nutr. 55, 349–360.

Dimidi, E., Cox, S.R., Rossi, M., and Whelan, K. (2019). Fermented Foods: Definitions and Characteristics, Impact on the Gut Microbiota and Effects on Gastrointestinal Health and Disease. Nutrients 11.

Duncan, S.H., Belenguer, A., Holtrop, G., Johnstone, A.M., Flint, H.J., and Lobley, G.E. (2007). Reduced dietary intake of carbohydrates by obese subjects results in decreased concentrations of butyrate and butyrate-producing bacteria in feces. Appl. Environ. Microbiol. 73, 1073–1078.

Earle, K.A., Billings, G., Sigal, M., Lichtman, J.S., Hansson, G.C., Elias, J.E., Amieva, M.R., Huang, K.C., and Sonnenburg, J.L. (2015). Quantitative Imaging of Gut Microbiota Spatial Organization. Cell Host Microbe 18, 478–488.

Fillatreau, S. (2018). B cells and their cytokine activities implications in human diseases. Clin. Immunol. Orlando Fla 186, 26–31.

Flint, H.J., Duncan, S.H., and Louis, P. (2017). The impact of nutrition on intestinal bacterial communities. Curr. Opin. Microbiol. 38, 59–65.

Fragiadakis, G.K., Bjornson-Hooper, Z.B., Madhireddy, D., Sachs, K., Spitzer, M.H., Bendall, S.C., and Nolan, G.P. (2019). Variation of immune cell responses in humans reveals sex-specific coordinated signaling across cell types (Immunology).

Fragiadakis, G.K., Wastyk, H.C., Robinson, J.L., Sonnenburg, E.D., Sonnenburg, J.L., and Gardner, C.D. (2020). Long-term dietary intervention reveals resilience of the gut microbiota despite changes in diet and weight. Am. J. Clin. Nutr. 111, 1127–1136.

GBD 2015 Mortality and Causes of Death Collaborators (2016). Global, regional, and national life expectancy, all-cause mortality, and cause-specific mortality for 249 causes of death, 1980-2015: a systematic analysis for the Global Burden of Disease Study 2015. Lancet Lond. Engl. 388, 1459–1544.

Gilbert, J.A., Jansson, J.K., and Knight, R. (2014). The Earth Microbiome project: successes and aspirations. BMC Biol. 12, 69.

Gille, D., Schmid, A., Walther, B., and Vergères, G. (2018). Fermented Food and Non-Communicable Chronic Diseases: A Review. Nutrients 10.

Gonzalez, C.G., Wastyk, H.C., Topf, M., Gardner, C.D., Sonnenburg, J.L., and Elias, J.E. (2020). High-Throughput Stool Metaproteomics: Method and Application to Human Specimens. MSystems 5, e00200–20, /msystems/5/3/msys.00200-20.atom.

Granado-Serrano, A.B., Martín-Garí, M., Sánchez, V., Riart Solans, M., Berdún, R., Ludwig, I.A., Rubió, L., Vilaprinyó, E., Portero-Otín, M., and Serrano, J.C.E. (2019). Faecal bacterial and short-chain fatty acids signature in hypercholesterolemia. Sci. Rep. 9, 1772.

Huang, L., Zhang, H., Wu, P., Entwistle, S., Li, X., Yohe, T., Yi, H., Yang, Z., and Yin, Y. (2018). dbCAN-seq: a database of carbohydrate-active enzyme (CAZyme) sequence and annotation. Nucleic Acids Res. 46, D516–D521.

Jha, A.R., Davenport, E.R., Gautam, Y., Bhandari, D., Tandukar, S., Ng, K.M., Fragiadakis, G.K., Holmes, S., Gautam, G.P., Leach, J., et al. (2018). Gut microbiome transition across a lifestyle gradient in Himalaya. PLoS Biol. 16, e2005396.

Johnson, A.J., Vangay, P., Al-Ghalith, G.A., Hillmann, B.M., Ward, T.L., Shields-Cutler, R.R., Kim, A.D., Shmagel, A.K., Syed, A.N., Walter, J., et al. (2019). Daily Sampling Reveals Personalized Diet-Microbiome Associations in Humans. Cell Host Microbe 25, 789–802.e5.

Klimenko, N.S., Tyakht, A.V., Popenko, A.S., Vasiliev, A.S., Altukhov, I.A., Ischenko, D.S., Shashkova, T.I., Efimova, D.A., Nikogosov, D.A., Osipenko, D.A., et al. (2018). Microbiome Responses to an Uncontrolled Short-Term Diet Intervention in the Frame of the Citizen Science Project. Nutrients 10.

Le Chatelier, E., Nielsen, T., Qin, J., Prifti, E., Hildebrand, F., Falony, G., Almeida, M., Arumugam, M., Batto, J.-M., Kennedy, S., et al. (2013). Richness of human gut microbiome correlates with metabolic markers. Nature 500, 541–546.

Lin, D., Peters, B.A., Friedlander, C., Freiman, H.J., Goedert, J.J., Sinha, R., Miller, G., Bernstein, M.A., Hayes, R.B., and Ahn, J. (2018). Association of dietary fibre intake and gut microbiota in adults. Br. J. Nutr. 120, 1014–1022.

Liu, L., Wang, S., and Liu, J. (2015). Fiber consumption and all-cause, cardiovascular, and cancer mortalities: a systematic review and meta-analysis of cohort studies. Mol. Nutr. Food Res. 59, 139–146.

Liu, S., Li, E., Sun, Z., Fu, D., Duan, G., Jiang, M., Yu, Y., Mei, L., Yang, P., Tang, Y., et al. (2019). Altered gut microbiota and short chain fatty acids in Chinese children with autism spectrum disorder. Sci. Rep. 9, 287.

Lozano, R., Naghavi, M., Foreman, K., Lim, S., Shibuya, K., Aboyans, V., Abraham, J., Adair, T., Aggarwal, R., Ahn, S.Y., et al. (2012). Global and regional mortality from 235 causes of death for 20 age groups in 1990 and 2010: a systematic analysis for the Global Burden of Disease Study 2010. Lancet Lond. Engl. 380, 2095–2128.

Lynch, S.V., and Pedersen, O. (2016). The Human Intestinal Microbiome in Health and Disease. N. Engl. J. Med. 375, 2369–2379.

Makki, K., Deehan, E.C., Walter, J., and Bäckhed, F. (2018). The Impact of Dietary Fiber on Gut Microbiota in Host Health and Disease. Cell Host Microbe 23, 705–715.

Martínez, I., Lattimer, J.M., Hubach, K.L., Case, J.A., Yang, J., Weber, C.G., Louk, J.A., Rose, D.J., Kyureghian, G., Peterson, D.A., et al. (2013). Gut microbiome composition is linked to whole grain-induced immunological improvements. ISME J. 7, 269–280.

Martínez, I., Stegen, J.C., Maldonado-Gómez, M.X., Eren, A.M., Siba, P.M., Greenhill, A.R., and Walter, J. (2015). The gut microbiota of rural papua new guineans: composition, diversity patterns, and ecological processes. Cell Rep. 11, 527–538.

Mosca, A., Leclerc, M., and Hugot, J.P. (2016). Gut Microbiota Diversity and Human Diseases: Should We Reintroduce Key Predators in Our Ecosystem? Front. Microbiol. 7, 455.

Mozaffarian, D., Hao, T., Rimm, E.B., Willett, W.C., and Hu, F.B. (2011). Changes in diet and lifestyle and long-term weight gain in women and men. N. Engl. J. Med. 364, 2392–2404.

Muegge, B.D., Kuczynski, J., Knights, D., Clemente, J.C., González, A., Fontana, L., Henrissat, B., Knight, R., and Gordon, J.I. (2011). Diet drives convergence in gut microbiome functions across mammalian phylogeny and within humans. Science 332, 970–974.

O’Keefe, S.J.D., Li, J.V., Lahti, L., Ou, J., Carbonero, F., Mohammed, K., Posma, J.M., Kinross, J., Wahl, E., Ruder, E., et al. (2015). Fat, fibre and cancer risk in African Americans and rural Africans. Nat. Commun. 6, 6342.

Ran-Ressler, R.R., Bae, S., Lawrence, P., Wang, D.H., and Brenna, J.T. (2014). Branched-chain fatty acid content of foods and estimated intake in the USA. Br. J. Nutr. 112, 565–572.

Rho, M., Tang, H., and Ye, Y. (2010). FragGeneScan: predicting genes in short and error-prone reads. Nucleic Acids Res. 38, e191–e191.

Rothschild, D., Weissbrod, O., Barkan, E., Kurilshikov, A., Korem, T., Zeevi, D., Costea, P.I., Godneva, A., Kalka, I.N., Bar, N., et al. (2018). Environment dominates over host genetics in shaping human gut microbiota. Nature 555, 210–215.

Smits, S.A., Leach, J., Sonnenburg, E.D., Gonzalez, C.G., Lichtman, J.S., Reid, G., Knight, R., Manjurano, A., Changalucha, J., Elias, J.E., et al. (2017). Seasonal cycling in the gut microbiome of the Hadza hunter-gatherers of Tanzania. Science 357, 802–806.

So, D., Whelan, K., Rossi, M., Morrison, M., Holtmann, G., Kelly, J.T., Shanahan, E.R., Staudacher, H.M., and Campbell, K.L. (2018). Dietary fiber intervention on gut microbiota composition in healthy adults: a systematic review and meta-analysis. Am. J. Clin. Nutr. 107, 965–983.

Sonnenburg, E.D., and Sonnenburg, J.L. (2014). Starving our microbial self: the deleterious consequences of a diet deficient in microbiota-accessible carbohydrates. Cell Metab. 20, 779–786.

Sonnenburg, E.D., and Sonnenburg, J.L. (2019). The ancestral and industrialized gut microbiota and implications for human health. Nat. Rev. Microbiol. 17, 383–390.

Sonnenburg, E.D., Smits, S.A., Tikhonov, M., Higginbottom, S.K., Wingreen, N.S., and Sonnenburg, J.L. (2016). Diet-induced extinctions in the gut microbiota compound over generations. Nature 529, 212–215.

Spencer, S.P., Fragiadakis, G.K., and Sonnenburg, J.L. (2019). Pursuing Human-Relevant Gut Microbiota-Immune Interactions. Immunity 51, 225–239.

Svedlund, J., Sjödin, I., and Dotevall, G. (1988). GSRS--a clinical rating scale for gastrointestinal symptoms in patients with irritable bowel syndrome and peptic ulcer disease. Dig. Dis. Sci. 33, 129–134.

Tanaka, T., Narazaki, M., and Kishimoto, T. (2014). IL-6 in inflammation, immunity, and disease. Cold Spring Harb. Perspect. Biol. 6, a016295.

Taylor, B.C., Lejzerowicz, F., Poirel, M., Shaffer, J.P., Jiang, L., Aksenov, A., Litwin, N., Humphrey, G., Martino, C., Miller-Montgomery, S., et al. (2020). Consumption of Fermented Foods Is Associated with Systematic Differences in the Gut Microbiome and Metabolome. MSystems 5.

The Human Microbiome Project Consortium (2012). A framework for human microbiome research. Nature 486, 215–221.

Turnbaugh, P.J., Hamady, M., Yatsunenko, T., Cantarel, B.L., Duncan, A., Ley, R.E., Sogin, M.L., Jones, W.J., Roe, B.A., Affourtit, J.P., et al. (2009). A core gut microbiome in obese and lean twins. Nature 457, 480–484.

Valles-Colomer, M., Falony, G., Darzi, Y., Tigchelaar, E.F., Wang, J., Tito, R.Y., Schiweck, C., Kurilshikov, A., Joossens, M., Wijmenga, C., et al. (2019). The neuroactive potential of the human gut microbiota in quality of life and depression. Nat. Microbiol. 4, 623–632.

Vangay, P., Johnson, A.J., Ward, T.L., Al-Ghalith, G.A., Shields-Cutler, R.R., Hillmann, B.M., Lucas, S.K., Beura, L.K., Thompson, E.A., Till, L.M., et al. (2018). US Immigration Westernizes the Human Gut Microbiome. Cell 175, 962–972.e10.

Walker, A.W., Ince, J., Duncan, S.H., Webster, L.M., Holtrop, G., Ze, X., Brown, D., Stares, M.D., Scott, P., Bergerat, A., et al. (2011). Dominant and diet-responsive groups of bacteria within the human colonic microbiota. ISME J. 5, 220–230.

Winham, D.M., and Hutchins, A.M. (2011). Perceptions of flatulence from bean consumption among adults in 3 feeding studies. Nutr. J. 10, 128.

Wu, G.D., Chen, J., Hoffmann, C., Bittinger, K., Chen, Y.-Y., Keilbaugh, S.A., Bewtra, M., Knights, D., Walters, W.A., Knight, R., et al. (2011). Linking long-term dietary patterns with gut microbial enterotypes. Science 334, 105–108.

Yatsunenko, T., Rey, F.E., Manary, M.J., Trehan, I., Dominguez-Bello, M.G., Contreras, M., Magris, M., Hidalgo, G., Baldassano, R.N., Anokhin, A.P., et al. (2012). Human gut microbiome viewed across age and geography. Nature 486, 222–227.

Yin, Y., Mao, X., Yang, J., Chen, X., Mao, F., and Xu, Y. (2012). dbCAN: a web resource for automated carbohydrate-active enzyme annotation. Nucleic Acids Res. 40, W445–451.

Zeevi, D., Korem, T., Zmora, N., Israeli, D., Rothschild, D., Weinberger, A., Ben-Yacov, O., Lador, D., Avnit-Sagi, T., Lotan-Pompan, M., et al. (2015). Personalized Nutrition by Prediction of Glycemic Responses. Cell 163, 1079–1094.

Zhernakova, A., Kurilshikov, A., Bonder, M.J., Tigchelaar, E.F., Schirmer, M., Vatanen, T., Mujagic, Z., Vila, A.V., Falony, G., Vieira-Silva, S., et al. (2016). Population-based metagenomics analysis reveals markers for gut microbiome composition and diversity. Science 352, 565–569.

(2014). Butyrate: food sources, functions and health benefits (New York: Nova Science Publishers, Inc).

