## Supplemental_figures_tables for "Gut Microbiota-Targeted Diets Modulate Human Immune Status"

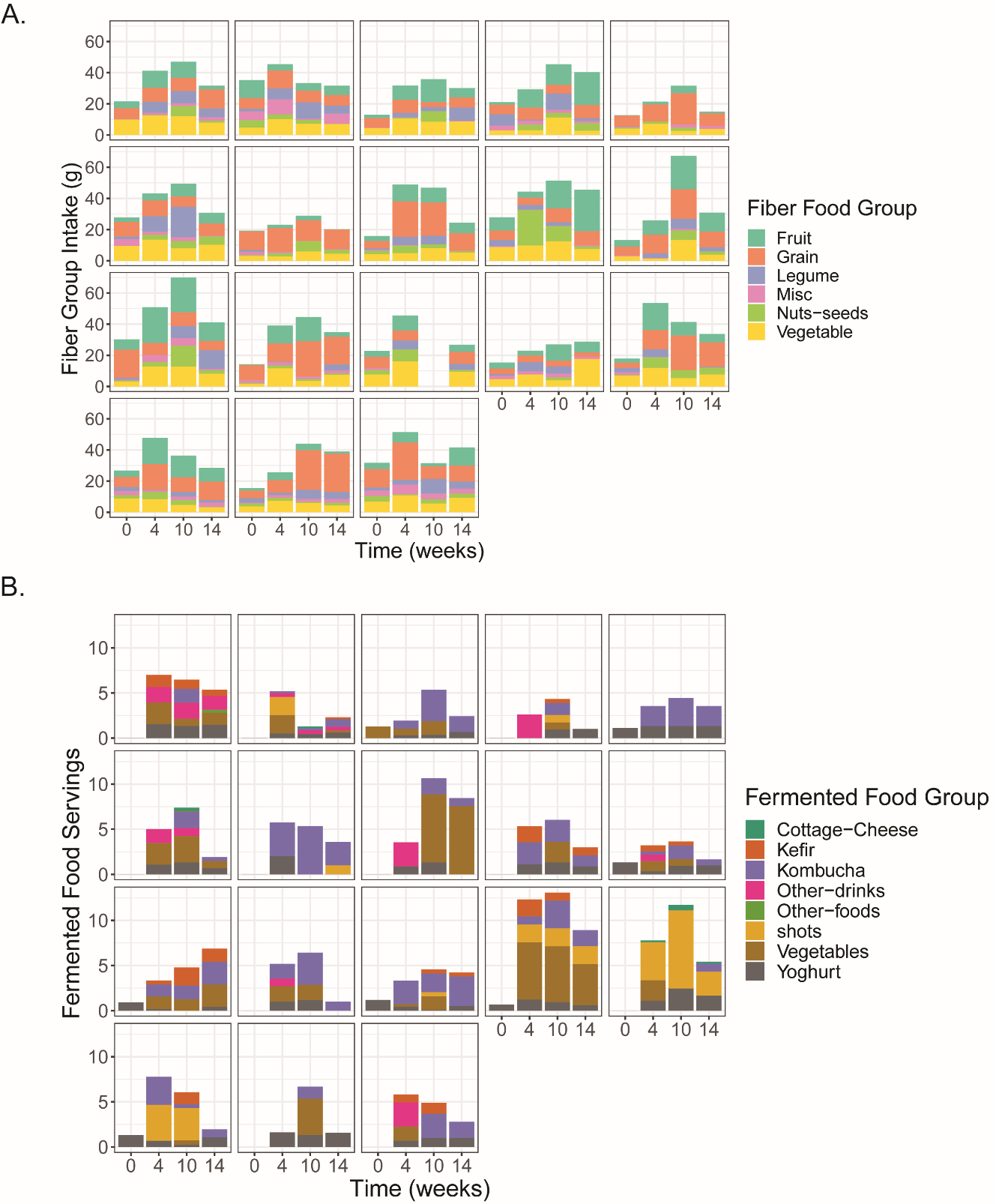
**Figure S1, Related to Figure 1. Individual participant high-fiber and high-fermented food diet arm group intake**.

(**A)** Participant-specific fiber intake by category for high-fiber diet arm.

(**B)** Participant-specific intake by category for high-fermented foods diet arm.

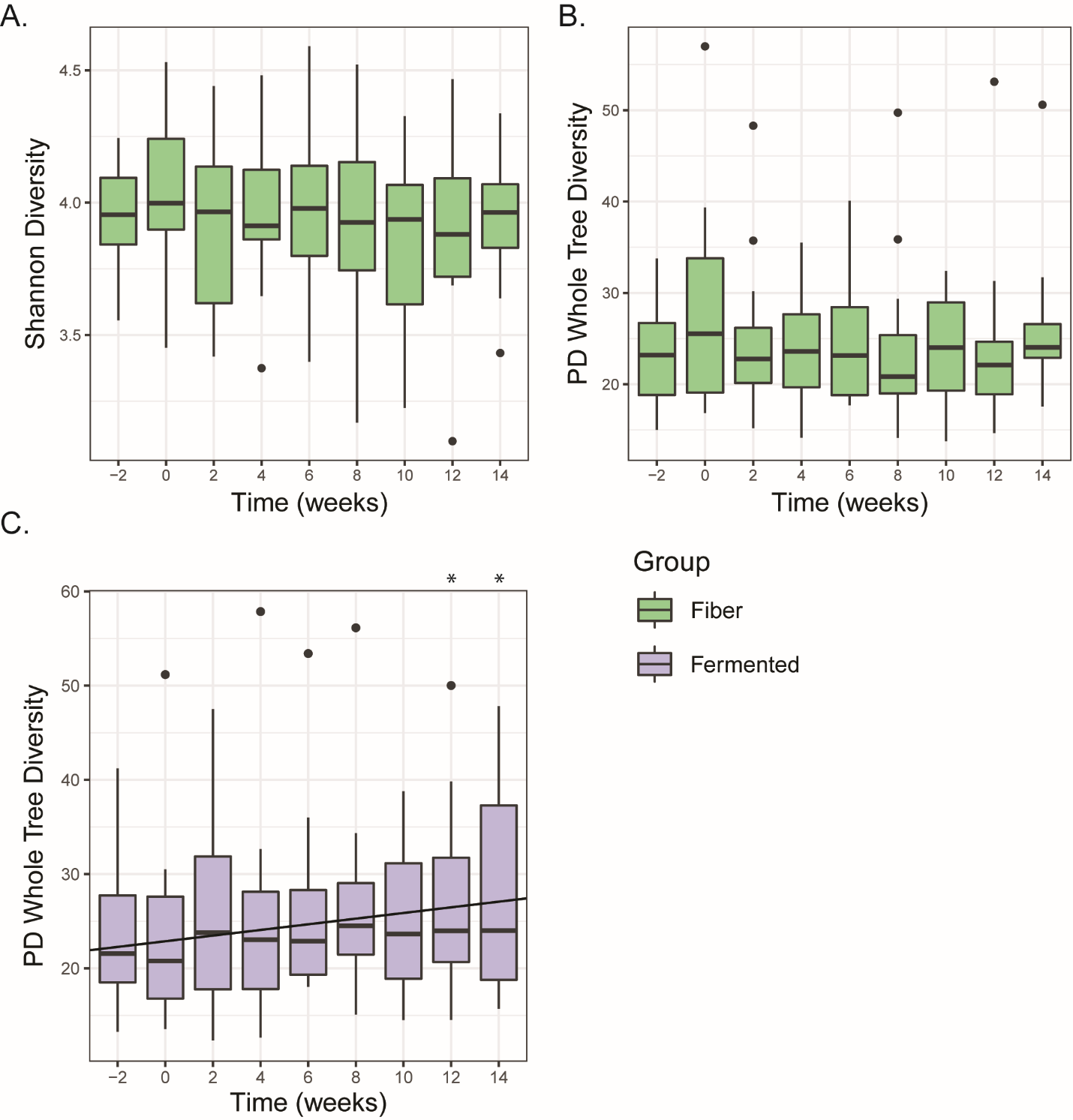

**Figure S2, Related to Figure 3 and 5. Alpha diversity measures for high-fiber and high-fermented diet arms**.

(**A)** Shannon alpha diversity in high-fiber diet arm.

(**B)** Phylogenetic Diversity **(**PD) whole tree alpha diversity in high-fiber food diet arm.

(**C)** Phylogenetic Diversity (PD) whole tree alpha diversity in high-fermented food diet arm. * indicates significant p-value relative to Week -2 (p-value < 0.05). Linear regression significant using a linear mixed effects model (p-value=0.013).

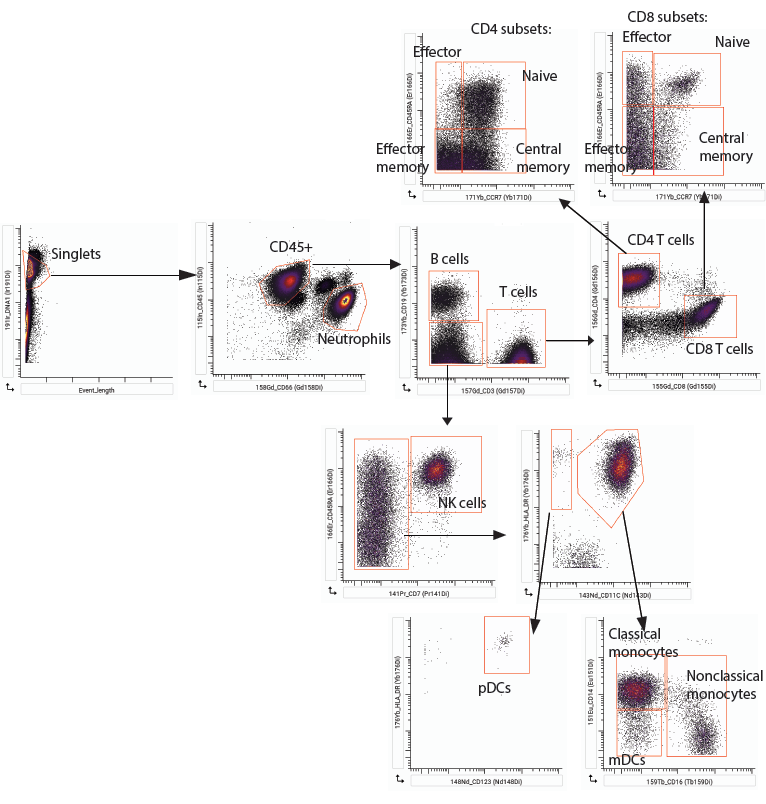

**Figure S3, Related to Figure 4 & 6. CyTOF gating strategy**.

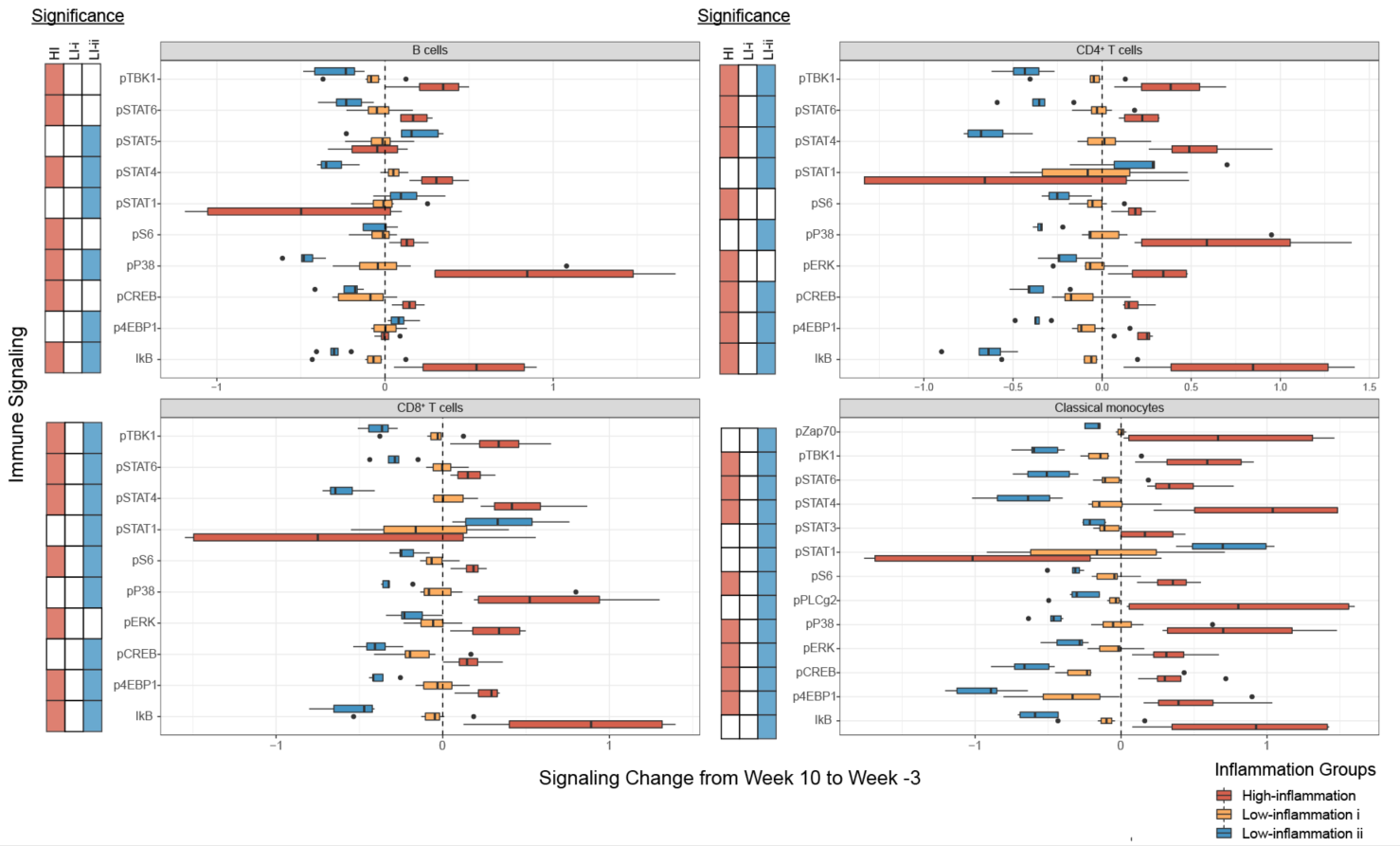

**Figure S4, Related to Figure 4. Changes in endogenous signaling in the high-fiber diet arm inflammation groups.** Endogenous signaling levels measured by CyTOF and identified as significantly changed (FDR < 0.05, q-value < 0.1, SAM test using siggenes package) from Baseline (Week -3) to end of Maintenance (Week 10) indicated by the shaded boxes next to the boxplots (Red=significantly changed in High-inflammation group, Blue=significantly changed in Low-inflammation ii group).

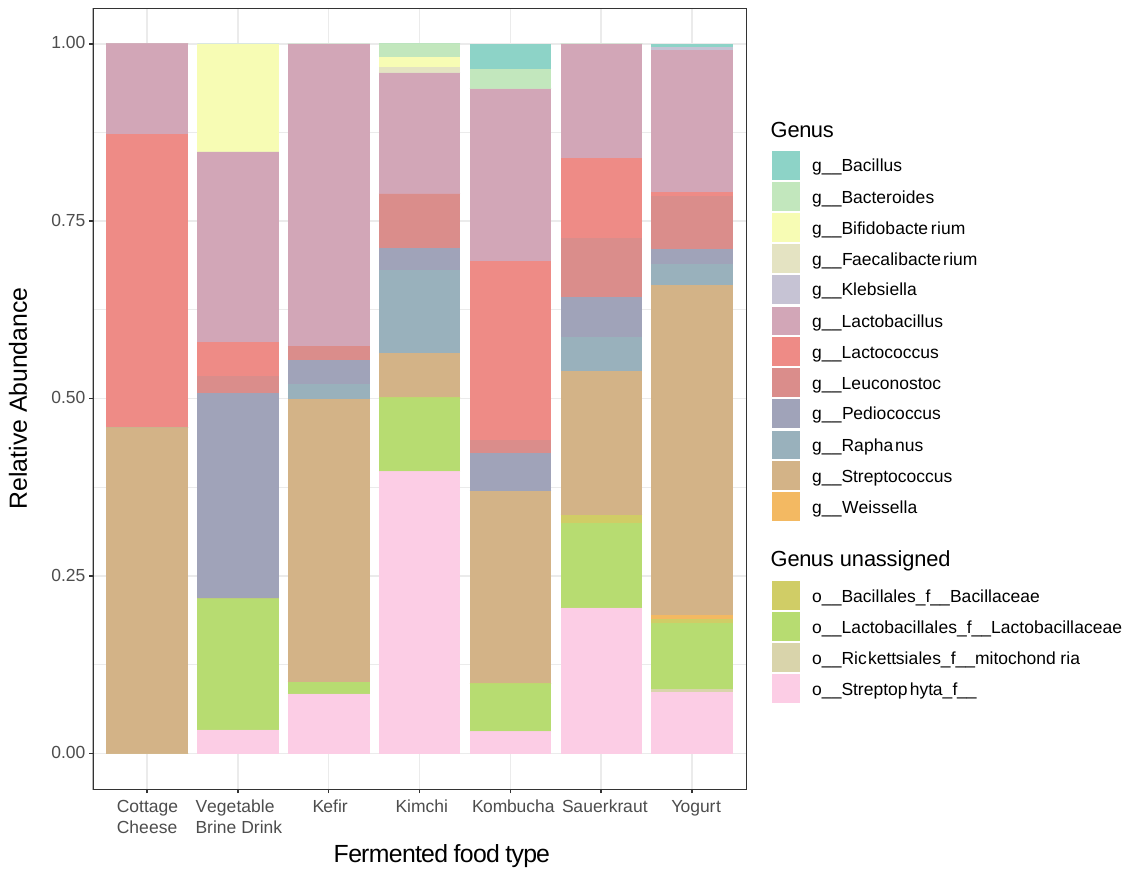

**Figure S5, Related to Figure 5. 16S ASV analysis of fermented foods.** ASVs with fewer than 250 counts were filtered out and counts binning to the same genus were summed. If genus was unassigned, ASVs were summed based on the same order and family.

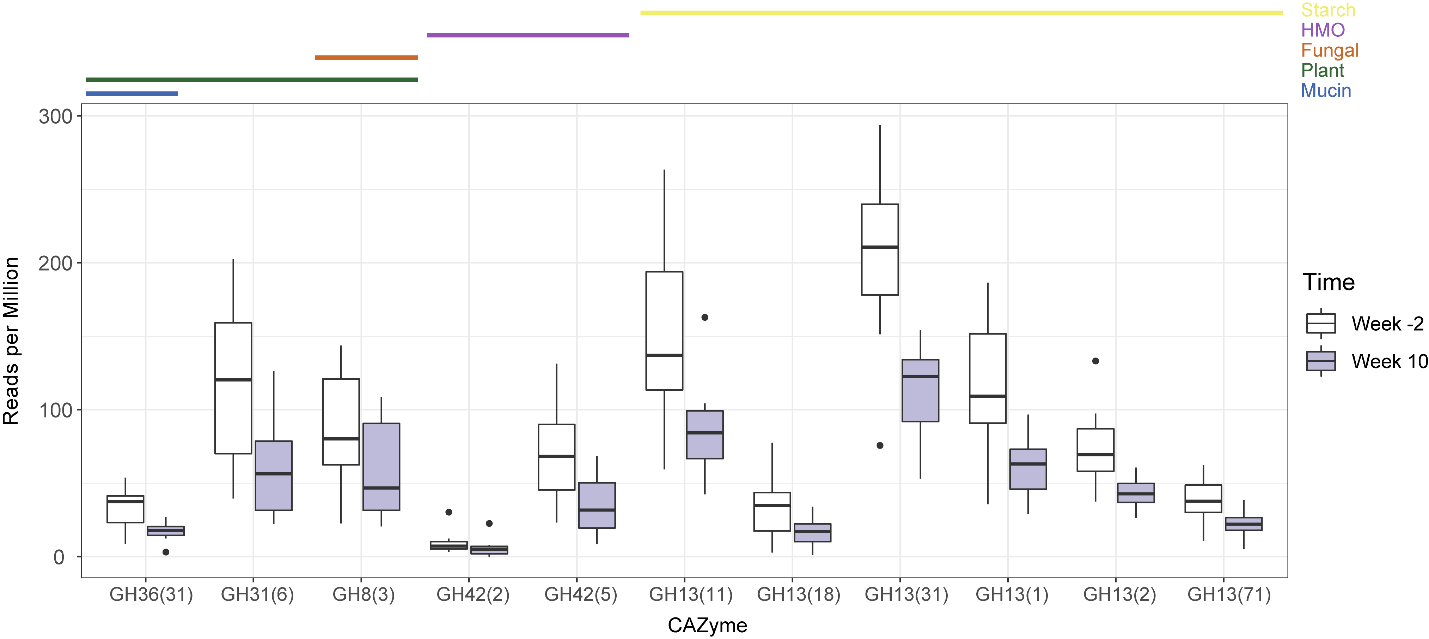

**Figure S6, Related to Figure 5. CAZyme analysis of high-fermented food diet arm.** CAZymes identified from metagenomic sequencing as significantly changing in relative abundance from baseline to end of maintenance phase (FDF ≤ 0.05, q-value ≤ 0.1, SAM test using siggenes package). CAZymes were annotated using dbCan and assigned to functional categories.

### Supplemental Tables

#### Table S1, Related to Figure 1. Demographics table.

|  | High fermented food diet (n=18) | High fiber diet (n=18) |
| --- | --- | --- |
| **Demographics** | | |
| Female | 13 | 12 |
| Male | 5 | 6 |
| Non-hispanic | 18 | 18 |
| Hispanic | 0 | 0 |
| Asian | 4 | 2 |
| White | 14 | 16 |
| Married/Partnered | 9 | 11 |
| Divorced | 7 | 1 |
| Single/Never married | 2 | 6 |
| Some college | 2 | 0 |
| College graduate | 5 | 6 |
| Some post-graduate school | 2 | 3 |
| Post-graduate degree | 9 | 9 |
| Working full-time | 14 | 15 |
| Working part-time | 1 | 2 |
| Unemployed | 0 | 1 |
| Retired | 3 | 0 |
| Smokers | 0 | 0 |
| Non-smokers | 18 | 16 |
| **Anthropometrics (average)** | | |
| HEIGHT (m) | 1.7 | 1.7 |
| WEIGHT (kg) | 70 | 73 |
| BMI | 25 | 25 |
| WAIST (cm) | 35 | 35 |
| Systolic BP | 121 | 120 |
| Diastolic BP | 77 | 76 |
| **Blood Values (average)** | | |
| Glucose (mg/dL) | 95 | 95 |
| Insulin (uU/mL) | 9.9 | 9.4 |
| Triglycerides (mg/dL) | 112 | 108 |
| Total Cholesterol (mg/dL) | 215 | 200 |
| HDL Cholesterol (mg/dL) | 65 | 61 |
| Calculated LDL Chol (mg/dL) | 128 | 117 |

#### Table S2, Related to Figure 1. Number of participants per data type.

|  | **Time (weeks)** | | | | | | | | | |
| --- | --- | --- | --- | --- | --- | --- | --- | --- | --- | --- |
|  | **Baseline** | | | **Ramp** | | **Maintenance** | | | **Choice** | |
|  | **-3** | **-2** | **0** | **2** | **4** | **6** | **8** | **10** | **12** | **14** |
| **Food logs** | 0 | 36 | 36 | 36 | 36 | 36 | 36 | 36 | 36 | 36 |
| **16S ASVs** | 0 | 32 | 28 | 28 | 31 | 33 | 33 | 33 | 32 | 31 |
| **Metagenomics** | 0 | 27 | 30 | 0 | 0 | 0 | 29 | 28 | 0 | 0 |
| **Proteomics** | 0 | 27 | 27 | 27 | 0 | 0 | 27 | 27 | 0 | 0 |
| **SCFAs** | 0 | 36 | 36 | 0 | 0 | 0 | 36 | 36 | 0 | 0 |
| **Inflammatory cytokines** | 35 | 0 | 35 | 0 | 0 | 33 | 35 | 35 | 0 | 0 |
| **Endogenous cell signaling** | 35 | 0 | 30 | 0 | 0 | 28 | 34 | 33 | 0 | 0 |
| **Cell signaling capacity** | 36 | 0 | 36 | 0 | 0 | 0 | 36 | 36 | 0 | 0 |

#### Table S3, Related to Figure 1. Nutrient data from participant diets.

##### A) High-fiber diet arm nutrient intake

| **Nutrient** | **Baseline** | **End of Maintenance** | **adjusted q-value** |
| --- | --- | --- | --- |
| Magnesium (mg) | 341.2+/-104.8 | 462.4+/-150.7 | **4.59E-04** |
| Potassium (mg) | 2726.4+/-549.1 | 3377+/-944.3 | **4.59E-04** |
| Vitamin C (ascorbic acid) (mg) | 77.3+/-54.9 | 166+/-118 | **4.59E-04** |
| Vitamin K (phylloquinone) (mcg) | 166.9+/-106.2 | 308.1+/-188.6 | **4.59E-04** |
| Total Dietary Fiber (g) | 22+/-7.4 | 43.3+/-15 | **2.49E-03** |
| Insoluble Dietary Fiber (g) | 15.7+/-5.6 | 33.1+/-11.9 | **2.86E-03** |
| Sodium (mg) | 2989.7+/-691.4 | 2603.1+/-1011.5 | **2.86E-03** |
| Animal Protein (g) | 49.1+/-17 | 32.9+/-15.4 | **3.67E-03** |
| Beta-Carotene (provitamin A carotenoid) (mcg) | 4574.8+/-3743.2 | 6532.1+/-4729.9 | **5.94E-03** |
| Total Sugars (g) | 79.7+/-28.8 | 97.6+/-35.8 | **6.14E-03** |
| Vegetable Protein (g) | 29.8+/-6.4 | 40.9+/-14.2 | **7.77E-03** |
| Calcium (mg) | 949.9+/-286 | 1026.7+/-275.5 | **7.92E-03** |
| Total Carbohydrate (g) | 224+/-58.1 | 248.1+/-63.3 | **8.11E-03** |
| Lutein + Zeaxanthin (mcg) | 3580.9+/-4969.5 | 4590.6+/-3312.1 | **1.10E-02** |
| Iron (mg) | 13.7+/-4.6 | 19+/-5.6 | **1.13E-02** |
| Energy (kcal) | 1880.4+/-366.2 | 1952.9+/-462.4 | **1.16E-02** |
| Alpha-Carotene (provitamin A carotenoid) (mcg) | 808.2+/-923.5 | 1009.6+/-1353 | **1.17E-02** |
| Soluble Dietary Fiber (g) | 6.3+/-2.4 | 10.1+/-4.8 | **1.17E-02** |
| Added Sugars (by Total Sugars) (g) | 36.7+/-23.9 | 37.8+/-25.3 | >.05 |
| Beta-Cryptoxanthin (provitamin A carotenoid) (mcg) | 354.4+/-728.1 | 296.1+/-348.3 | >.05 |
| Cholesterol (mg) | 224.9+/-61.8 | 196.3+/-131.5 | >.05 |
| Lycopene (mcg) | 3992.5+/-4108.2 | 3720.7+/-4969.1 | >.05 |
| Pectins (g) | 2.8+/-1.3 | 5.8+/-2.8 | >.05 |
| Refined Grains (ounce equivalents) | 4.8+/-2.5 | 2.9+/-1.1 | >.05 |
| Total Fat (g) | 71+/-17.7 | 77.5+/-25.9 | >.05 |
| Total Grains (ounce equivalents) | 7.4+/-3 | 6+/-2.2 | >.05 |
| Total Monounsaturated Fatty Acids (MUFA) (g) | 24.5+/-6.6 | 27.9+/-11.6 | >.05 |
| Total Polyunsaturated Fatty Acids (PUFA) (g) | 16.4+/-5.2 | 19.7+/-6.6 | >.05 |
| Total Protein (g) | 78.9+/-18.6 | 73.9+/-20.1 | >.05 |
| Total Saturated Fatty Acids (SFA) (g) | 23.8+/-7.4 | 23.2+/-10.7 | >.05 |
| Whole Grains (ounce equivalents) | 2.6+/-3.6 | 3.1+/-2 | >.05 |

##### B) High-fermented diet arm nutrient intake

| **Nutrient** | **Baseline** | **End of Maintenance** | **adjusted q-value** |
| --- | --- | --- | --- |
| Animal Protein (g) | 45.4+/-18.9 | 58.3+/-20.1 | **1.49E-02** |
| Added Sugars (by Total Sugars) (g) | 37.7+/-17.5 | 50+/-19.1 | > .05 |
| Alpha-Carotene (provitamin A carotenoid) (mcg) | 632.3+/-785.3 | 456.4+/-566 | > .05 |
| Beta-Carotene (provitamin A carotenoid) (mcg) | 4876.1+/-3673.7 | 4465.6+/-2331.8 | > .05 |
| Beta-Cryptoxanthin (provitamin A carotenoid) (mcg) | 383.4+/-737.4 | 345.3+/-684.8 | > .05 |
| Calcium (mg) | 783.7+/-253.5 | 927.5+/-283.8 | > .05 |
| Cholesterol (mg) | 238.5+/-134.8 | 252.7+/-128.2 | > .05 |
| Energy (kcal) | 1801.1+/-437.9 | 1656.8+/-292.9 | > .05 |
| Insoluble Dietary Fiber (g) | 13.9+/-5.1 | 14.2+/-7.1 | > .05 |
| Iron (mg) | 11.6+/-2.4 | 10.8+/-2.9 | > .05 |
| Lutein + Zeaxanthin (mcg) | 2281.9+/-1873.4 | 3868.5+/-3240 | > .05 |
| Lycopene (mcg) | 2072.3+/-2152.5 | 1976.1+/-3023.9 | > .05 |
| Magnesium (mg) | 306.9+/-99.1 | 320.2+/-98.2 | > .05 |
| Pectins (g) | 2.9+/-1 | 3.4+/-1.5 | > .05 |
| Potassium (mg) | 2602.6+/-618.3 | 2822.8+/-803.9 | > .05 |
| Refined Grains (ounce equivalents) | 4.5+/-2.5 | 2.5+/-1.7 | > .05 |
| Sodium (mg) | 2534.6+/-798.2 | 2532.8+/-557.2 | > .05 |
| Soluble Dietary Fiber (g) | 6.7+/-2.4 | 6.2+/-2.7 | > .05 |
| Total Carbohydrate (g) | 205.3+/-54.3 | 188.6+/-46.9 | > .05 |
| Total Dietary Fiber (g) | 21.3+/-7.3 | 20.6+/-9.5 | > .05 |
| Total Fat (g) | 72.3+/-20.5 | 64.2+/-18 | > .05 |
| Total Grains (ounce equivalents) | 6.1+/-2 | 4+/-1.6 | > .05 |
| Total Monounsaturated Fatty Acids (MUFA) (g) | 27+/-7.4 | 23.4+/-6.6 | > .05 |
| Total Polyunsaturated Fatty Acids (PUFA) (g) | 17.4+/-6.4 | 14.1+/-6.9 | > .05 |
| Total Protein (g) | 74.1+/-21 | 81.2+/-18.2 | > .05 |
| Total Saturated Fatty Acids (SFA) (g) | 21.9+/-8.4 | 21.1+/-7.5 | > .05 |
| Total Sugars (g) | 78.7+/-27.4 | 93.2+/-28.3 | > .05 |
| Vegetable Protein (g) | 28.7+/-9.1 | 22.9+/-9.1 | > .05 |
| Vitamin C (ascorbic acid) (mg) | 92+/-60.2 | 107.1+/-51.7 | > .05 |
| Vitamin K (phylloquinone) (mcg) | 150+/-84.9 | 215.4+/-145.3 | > .05 |
| Whole Grains (ounce equivalents) | 1.6+/-1.6 | 1.5+/-1 | > .05 |

#### Table S4. Primary and Secondary Clinical Trial Outcomes.

Provided as Excel file: Table_S4_primary_secondary_outcomes.xlsx

#### Table S5, Related to Figure 2. Features selected in random forest models predicting diet group.

Provided as Excel file: Table_S5_loocv_opt_variables.xlsx

#### Table S6, Related to Figure 4 & 6. Immune profiling panels.

Provided as Excel file: Table_S6_immune_profiling_panels.xlsx.

Contains panels for Olink, CyTOF surface markers, CyTOF intracellular markers, and Phospho-flow

#### Table S7, Related to Figure 4. ZIBR coefficients for High-fiber diet arm inflammation clusters.

| Comparison | Phylogenetic assignment | | | Joint model | | Beta regression model | | | | Logarithmic model | | | |
| --- | --- | --- | --- | --- | --- | --- | --- | --- | --- | --- | --- | --- | --- |
|  |  |  |  | Adjusted p-value | | Group | | Baseline | | Group | | Baseline | |
|  | Family | Genus | Species | Group | Baseline | Coefficient | Adjusted p-val | Coefficient | Adjusted p-val | Coefficient | Adjusted p-val | Coefficient | Adjusted p-val |
| High v Low i | *Lachnospiraceae* | *Coprococcus* | UA | **0.01** | 0.07 | 0.31 | 0.82 | 0.38 | 1.00 | 3.09 | **0.01** | -2720.24 | 0.05 |
|  | *Ruminococcaceae* | *Oscillospira* | UA | 0.08 | 0.72 | -44.88 | 1.00 | 105.10 | 0.45 | 2.46 | **0.04** | -413.64 | 0.59 |
|  | *Ruminococcaceae* | *Ruminococcus* | UA | 0.11 | 0.25 | 0.03 | 1.00 | -10.74 | 0.92 | 1.56 | **0.04** | 264.09 | 0.16 |
|  | *Lachnospiraceae* | *Anaerostipes* | UA | **0.01** | 0.35 | -0.19 | 0.74 | -19.88 | 0.63 | 1.22 | 0.59 | 320.80 | 0.28 |
| High v Low ii | *Verrucomicrobiaceae* | *Akkermansia* | *muciniphila* | 0.20 | 1.00 | -1.35 | **0.04** | 0.26 | 1.00 | -6.03 | 0.40 | -2.71 | 1.00 |
| Low i v Low ii | *Lachnospiraceae* | UA | UA | 1.00 | 0.17 | -2.75 | **0.00** | 69.55 | 0.34 | 0.26 | 1.00 | 191.17 | 0.16 |

Note: UA = Unassigned taxa, significant p-values bolded.

ZIBR coefficients on the tip glommeration for fiber inflammation groups. Beta-regression model refers to the change in relative abundance of taxa. Logarithmic model refers to presence/absence of taxa. A significant coefficient for the grouping variable indicates a significant difference between the inflammation groups indicated in the comparison column. Baseline coefficients indicate a difference in the taxonomic relative abundance or presence/absence at the baseline timepoint (Week 0). Taxa with significant baseline coefficients were filtered out to focus on the significant differences induced by the dietary intervention.

#### Table S8, Related to Figure 7. Protein disease annotations.

Provided as Excel file: Table_S8_host_proteo_disease_annotation.xlsx
